# *Candida albicans’* inorganic phosphate transport and evolutionary adaptation to phosphate scarcity

**DOI:** 10.1101/2024.01.29.577887

**Authors:** Maikel Acosta-Zaldívar, Wanjun Qi, Abhishek Mishra, Udita Roy, William R. King, Jana Patton-Vogt, Matthew Z. Anderson, Julia R. Köhler

## Abstract

Phosphorus is essential in all cells’ structural, metabolic and regulatory functions. For fungal cells that import inorganic phosphate (Pi) up a steep concentration gradient, surface Pi transporters are critical capacitators of growth. Fungi must deploy Pi transporters that enable optimal Pi uptake in pH and Pi concentration ranges prevalent in their environments. Single, triple and quadruple mutants were used to characterize the four Pi transporters we identified for the human fungal pathogen *Candida albicans*, which must adapt to alkaline conditions during invasion of the host bloodstream and deep organs. A high-affinity Pi transporter, Pho84, was most efficient across the widest pH range while another, Pho89, showed high-affinity characteristics only within one pH unit of neutral. Two low-affinity Pi transporters, Pho87 and Fgr2, were active only in acidic conditions. Only Pho84 among the Pi transporters was clearly required in previously identified Pi-related functions including Target of Rapamycin Complex 1 signaling and hyphal growth. We used in vitro evolution and whole genome sequencing as an unbiased forward genetic approach to probe adaptation to prolonged Pi scarcity of two quadruple mutant lineages lacking all 4 Pi transporters. Lineage-specific genomic changes corresponded to divergent success of the two lineages in fitness recovery during Pi limitation. In this process, initial, large-scale genomic alterations like aneuploidies and loss of heterozygosity were eventually lost as populations presumably gained small-scale mutations. Severity of some phenotypes linked to Pi starvation, like cell wall stress hypersensitivity, decreased in parallel to evolving populations’ fitness recovery in Pi scarcity, while that of others like membrane stress responses diverged from these fitness phenotypes. *C. albicans* therefore has diverse options to reconfigure Pi management during prolonged scarcity. Since Pi homeostasis differs substantially between fungi and humans, adaptive processes to Pi deprivation may harbor small-molecule targets that impact fungal growth and virulence.

**Author Summary:** Fungi must be able to access enough phosphate in order to invade the human body. Virulence of *Candida albicans*, the most common invasive human fungal pathogen, is known to decrease when one of the proteins that brings phosphate into the fungal cell, called Pho84, is disabled. We identified three more proteins in *C. albicans* that transport phosphate into the cell. We found that Pho84 plays the largest role among them across the broadest range of environmental conditions. After eliminating all 4 of these transporters, we let two resulting mutants evolve for two months in limited phosphate and analyzed the growth and stress resistance of the resulting populations. We analyzed genomes of representative populations and found that early adaptations to phosphate scarcity occurred together with major changes to chromosome configurations. In later stages of the adaptation process, these large-scale changes disappeared as populations presumably gained small-scale mutations that increased cells’ ability to grow in limited phosphate. Some but not all of these favorable mutations improved resistance of evolving populations to stressors like membrane- and cell wall stress. Pinpointing distinct mutation combinations that affect stress resistance differently in populations adapting to scarce phosphate, may identify useful antifungal drug targets.

## Introduction

Phosphorus is an essential macronutrient for living cells and a major component of chromosomes, membranes and the transcription and translation machineries [1]. Inorganic phosphate (Pi) is required in the production of ATP, the energy currency of the cell, that governs central metabolic processes and intracellular signaling. Consequently, Pi is not only required for growth and proliferation but also for survival: e.g., fission yeast cells starved for Pi initially become quiescent and then lose viability [2].

Osmotrophic organisms that take up soluble small-molecule nutrients from their immediate environment must import Pi separately from molecular sources of nitrogen and carbon to acquire sufficient phosphorus. Most soils and aquatic environments contain <1%, or ≤10 mM soluble Pi, so that Pi is a scarce resource for plants and free-living microorganisms [3–5]; human serum Pi ranges from 0.8-1.3 mM [6], suggesting that microbial invasive pathogens of humans also experience Pi deprivation. For this reason, the Pi-homeostatic systems of small-molecule importing organisms like bacteria, plants and fungi have much in common. Orthology of phosphate proton symporters among plants and fungi was first determined by complementation of a *Saccharomyces cerevisiae* null mutant in *PHO84* with two *Arabidopsis thaliana* Pi transporters [7, 8]. In contrast, human phosphate homeostasis regulation differs fundamentally from that of osmotrophs [9, 10], and since abundant phosphorus-containing molecules are present in all human food sources of protein, the major high-affinity Pi transporter of fungi has no human homolog.

*Saccharomyces cerevisiae* Pi transporters were characterized over decades according to their Pi affinity and their pH optima [11]. Kinetic studies in the 1980s identified two Pi transport systems, one with a low K_m_ value of 8.4-21.4 µM, defined by the early investigators of these systems as high-affinity, and another with a high K_m_ value of 0.77-1.7 mM, defined as low-affinity [12, 13]. Further analysis suggested two separate transporters with distinct pH optima within the high-affinity uptake system [14]. Cloning and functional characterization of the *PHO84* gene showed that its product is a component of the high-affinity Pi transport system [15]. Heterologous expression of Pho84 and its incorporation into liposomes then permitted kinetic studies that demonstrated a K_m_ for Pi of 24 µM [16]. *S. cerevisiae* Pho84 is a member of the Phosphate: H^+^ Symporter Family within the Major Facilitator Superfamily [17]; it uses the chemiosmotic energy of proton symport to transport the Pi anion up a concentration gradient across the plasma membrane.

Additional Pi import systems were subsequently identified and characterized. A separate high-affinity Pi uptake system that was enhanced in the presence of sodium at pH 7.2 was described [18]; the gene encoding this activity was later identified and its product, named Pho89, confirmed to have a Pi K_m_ of 0.5 µM [14]. Pho89 belongs to solute carrier family 20 as a sodium-dependent phosphate transporter [19, 20]. Low-affinity *S. cerevisiae* Pi transporters Pho87, Pho90 and Pho91 were subsequently characterized genetically and functionally [21]. Pho91 was later shown to reside on the vacuolar membrane and facilitate Pi export from the vacuole to the cytosol [22]. Further work showed distinct activities of the 2 low-affinity transporters: Pho87 versus Pho90 can sustain growth of *S. cerevisiae* down to Pi concentrations of 5 mM versus 0.5 mM, respectively [23]. *S. cerevisiae* therefore has 2 high-affinity Pi transporters, Pho84 and Pho89, whose energetic drivers are proton- and sodium symport, respectively, and 2 paralogous low-affinity Pi transporters, Pho87 and Pho90 [21, 23].

The genome of the opportunistic fungal pathogen *Candida albicans* encodes 4 homologs of *S. cerevisiae* Pi transporters. In a *mariner* transposon mutant screen we previously identified a mutant in the *C. albicans* homolog of *S. cerevisiae PHO84* as hypersensitive to rapamycin [24]. We showed that *C. albicans PHO84* is required in normal Target of Rapamycin Complex 1 (TORC1) signaling, oxidative- and cell wall stress resistance, survival during exposure to amphotericin B and the echinocandin micafungin, and normal virulence [24–26]. Given the presence of other Pi transporter homologs in the *C. albicans* genome, we sought to understand how loss of just one, Pho84, could significantly impact important physiological functions and even virulence in *C. albicans*.

Pi-acquisition and -homeostatic systems (PHO regulons) of bacterial and other fungal human pathogens are required for virulence, implicating Pi scarcity as a prevalent condition in the host [27–30]: e.g., in a pathogen that is completely adapted to its human host, *Mycobacterium tuberculosis*, transcription of a secretion system for virulence factors is activated by Pi starvation [31, 32]. Expression of high-affinity Pi transporters is typically regulated according to ambient Pi concentrations [33]. In *S. cerevisiae*, high-affinity Pi transporter-encoding genes *PHO84* and *PHO89* are upregulated during Pi starvation [21, 34, 35]. In *C. albicans* ex vivo and in vivo infection models [36–40], the *PHO84* and *PHO89* homologs are similarly upregulated [14]. These findings suggest that in the host, *C. albicans* like *M. tuberculosis* experiences Pi starvation. We therefore set out to identify and characterize the other putative *C. albicans* Pi importers that can contribute to cytosolic Pi availability for the fungus’ growth and interaction with the host [24–26, 41].

We then asked whether *C. albicans* can adapt to persistent Pi scarcity. Forward genetic screens of chemically or transposon-mutagenized cells subjected to specific selective conditions are powerful discovery tools because they provide unbiased and often unexpected information [42]. In vitro evolution and whole genome analysis has been used in other pathogens for the characterization of drug responses [43–47] and in *Candida* species for analysis of drug resistance development [48–52]. We questioned whether this approach might have benefits for analysis of adaptation to nutrient scarcity compared with mutant screens. Its possible advantages might be a higher likelihood of revealing illuminating gain-of-function mutations and the ability to uncover mutations involved in polygenic traits. We used in vitro evolution and genome analysis to begin uncovering the cellular processes linked to *C. albicans’* management of Pi scarcity.

## Results

### *PHO84* plays a central role in growth and filamentation

Homology searches identified four Pi transporters in the *C. albicans* genome. A *C. albicans* ortholog of *S. cerevisiae PHO89,* which encodes a high-affinity Pi transporter with an alkaline optimum [53], resides on chromosome 4. Homology searches using the low affinity *S. cerevisiae PHO87* and *PHO90* paralogs found a single homolog named *PHO87* in the Candida Genome Database (CGD) [53]. A *PHO84* homolog named *FGR2* was also identified that shares 23% amino acid identity and 42% similarity with Pho84 (S1 Fig.)*. FGR2* was first isolated in a screen for *C. albicans* mutants impaired in filamentous growth [54] and more recently found to contribute to filamentation differences between *C. albicans* strains [55]. To delineate the contribution of these 4 predicted transporters to Pi acquisition, we constructed single gene deletion mutants for *PHO89, PHO87* and *FGR2* using our FLP-*NAT1* system [56] to compare with our previously constructed *pho84*-/- null mutants [24].

Pi restriction reduced growth only of *pho84*-/- mutants at acidic pH (Fig. 1). Null mutants in *PHO84* were unable to grow on synthetic complete (SC) agar medium containing low (0.05 mM) or moderate (0.5 mM) Pi at pH 3 or 5. In contrast, null mutants in the 3 other predicted Pi transporters grew similarly to the wildtype control strain (WT) under these conditions, indicating that only *PHO84* is required in moderate to low Pi at lower than neutral pH (Fig. 1; shown in Fig. 1 are also triple and quadruple mutants, which retain only one or none of the 4 Pi transporters, respectively; these are described below in detail). At pH 7, *pho84* null mutants grew at all tested Pi concentrations, indicating that one or more other Pi transporters were able to uptake sufficient Pi to sustain growth at neutral pH. At 7.3 mM Pi, all single mutants in the 4 predicted Pi transporters grew robustly, indicating that no single transporter is indispensable at high Pi concentrations (Fig. 1).

**Fig. 1.**
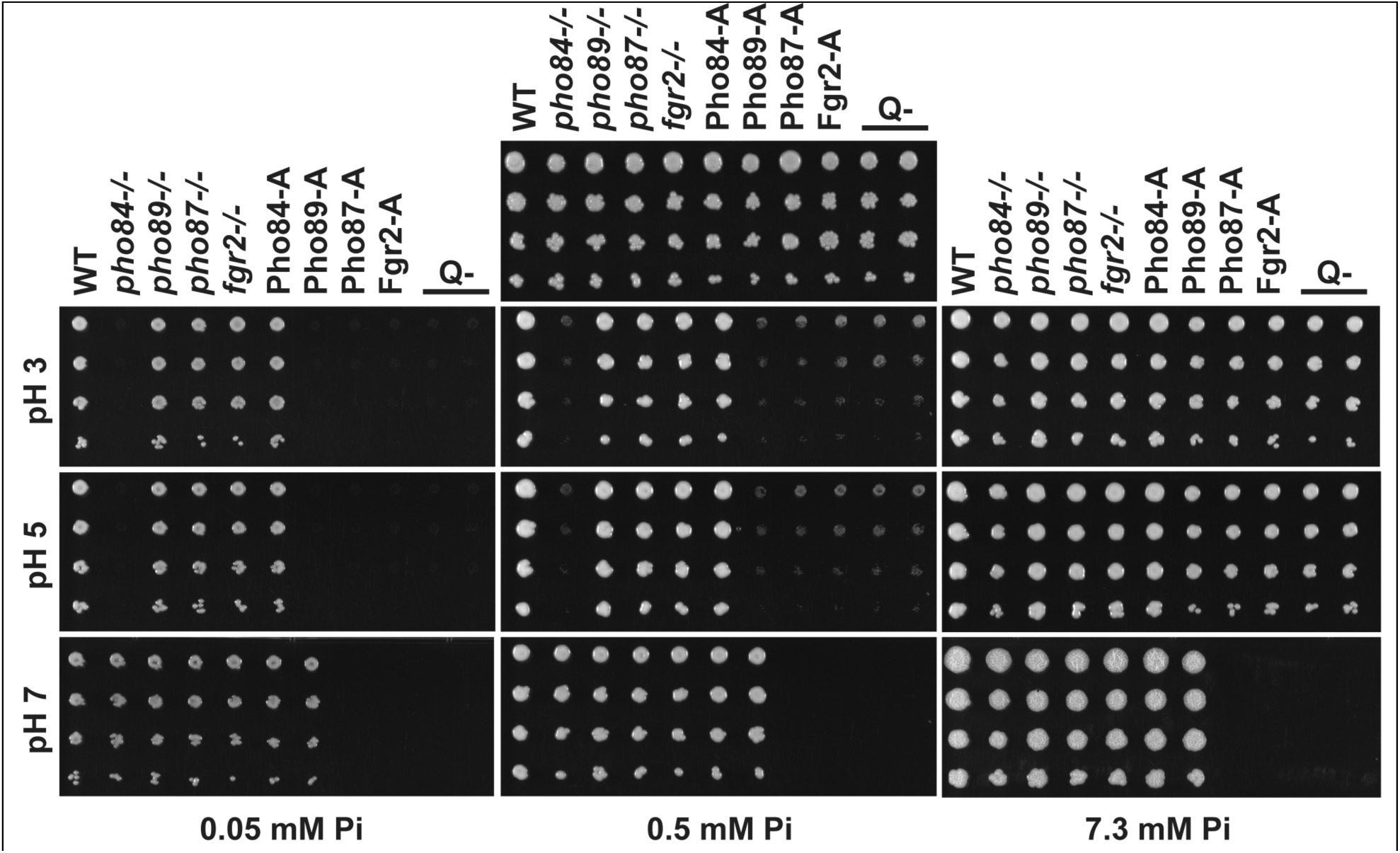
Among Pi transporters, Pho84 contributed to growth over conditions. Fivefold dilutions of cells of indicated genotypes were spotted (top to bottom) onto YPD (top, center) or SC (all others) agar media buffered to indicated pH (3, 5 or 7) with 100 mM MES and containing indicated Pi concentrations (0.05, 0.5 or 7.3 mM) and grown at 30° C for 2 d. Strains are WT (JKC915); *pho84-/-* (JKC1450); *pho89-/-* (JKC2585); *pho87-/-* (JKC2581); *fgr2-/-* (JKC2667); Pho84-A: *pho87-/- pho89- /- fgr2-/- PHO84+/+* (JKC2788); Pho89-A: *pho84-/- pho87-/- fgr2-/- PHO89+/+* (JKC2783); Pho87-A: *pho84-/- pho89-/- fgr2-/- PHO87+/+* (JKC2777); Fgr2-A: *pho84-/- pho87-/- pho89-/- FGR2+/+* (JKC2758); Q-: *pho84-/- pho87-/- pho89-/- fgr2-/-* (JKC2830 and JKC2860). Representative of 3 biological replicates.

*C. albicans’* ability to readily switch between growth as single budding yeast versus as multicellular filamentous hyphae contributes to its virulence [57, 58]. Cells lacking *PHO84* were previously shown to be defective in hyphal formation [25, 59]. To examine the role of the different Pi transporters in morphogenesis, cell suspensions from each of the 4 Pi transporter mutants were spotted on filamentation-inducing agar media. Most clearly on Spider medium, cells lacking *PHO84* had minimal or absent hyphal growth (Fig. 2A). Single mutants of the other 3 Pi transporters showed more subtle filamentation defects than *pho84*-/- cells. Similar filamentation phenotypes were observed on RPMI agar at pH 5 and pH 7: *pho84-/-* mutants produced occasional thin wisps of peripheral hyphae only on RPMI at pH 5 but not pH 7, while *pho89-/-*, *pho87-/-* and *fgr2-/-* mutants showed robust hyphal growth on these media (Fig. 2B). Together, these findings show that *PHO84* is the most important predicted Pi transporter for filamentation under these conditions.

**Fig. 2.**
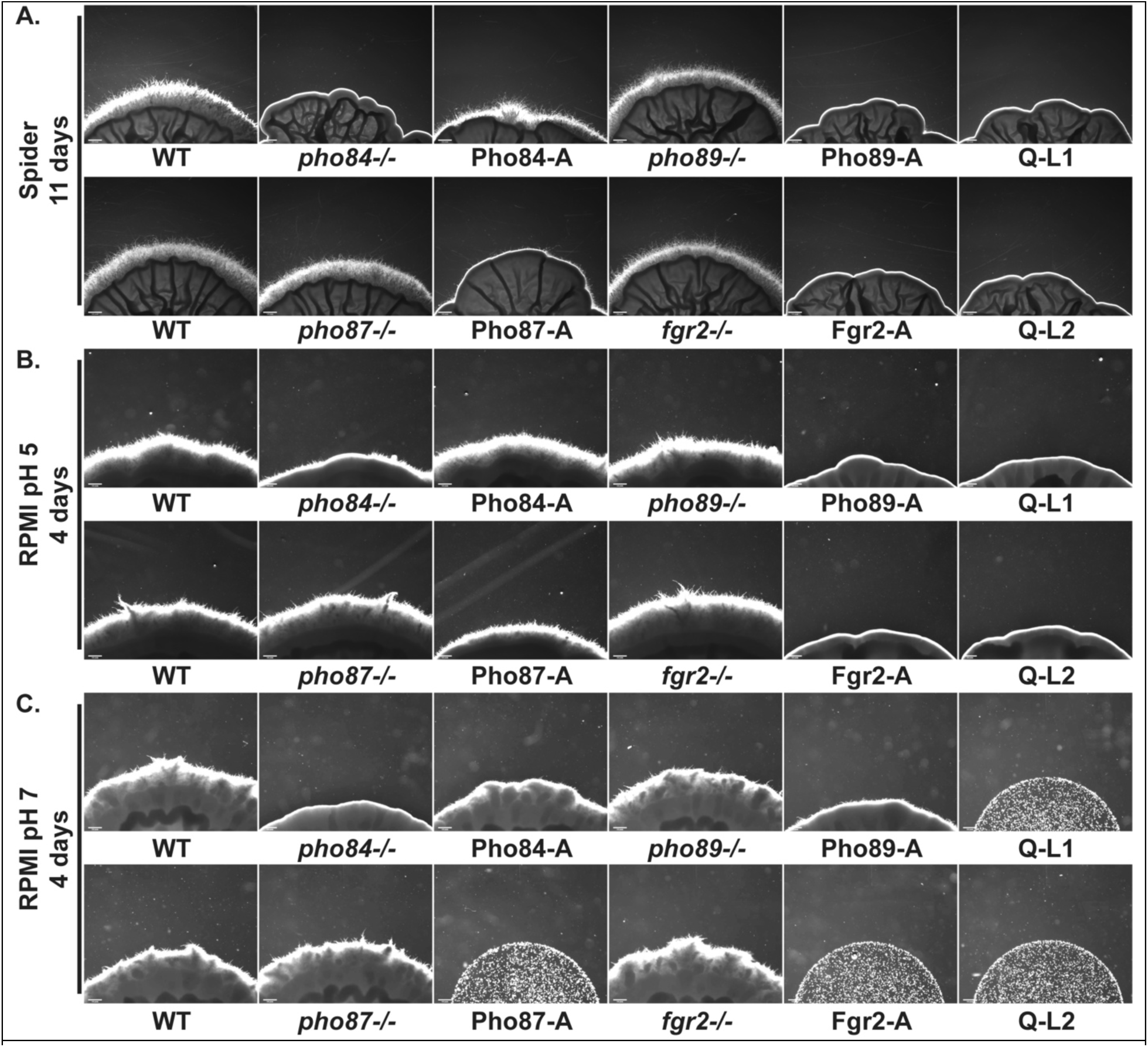
Pho84 was the major contributor to hyphal growth among 4 Pi transporters. Cell suspensions of indicated genotypes were spotted at equidistant points around the perimeter of Spider (**A**) and RPMI (**B, C**) agar plates. Photomicrographs of the edge of spots were obtained at 4 days for RPMI and 11 days for Spider plates. Spot edges were aligned with image frame corners to allow comparisons of hyphal fringes’ length. Strains are WT (JKC915); *pho84-/-* (JKC1450); Pho84-A: *pho87-/- pho89-/- fgr2-/- PHO84+/+* (JKC2788); *pho89-/-* (JKC2585); Pho89-A: *pho84-/- pho87-/- fgr2-/- PHO89+/+* (JKC2783)*; pho87-/-* (JKC2581); Pho87-A: *pho84-/- pho89-/- fgr2-/- PHO87+/+* (JKC2777); *fgr2-/-* (JKC2667); Fgr2-A: *pho84-/- pho87-/- pho89-/- FGR2+/+* (JKC2758); Q- L1: *pho84-/- pho87-/- pho89-/- fgr2-/-* (JKC2830); Q- L2: *pho84-/- pho87-/- pho89-/- fgr2-/-* (JKC2860). Size bar 200 µm. Representative of 3 biological replicates.

### Only *PHO84* impacted TORC1 signaling and oxidative stress endurance

We previously found that mutants in *PHO84* are hypersensitive to rapamycin and show decreased TORC1 signaling when growing in limited Pi [24], and that TORC1 co-regulates *PHO84* expression in addition to its known regulation by Pho4 [60, 61]. TORC1 activation was reduced in *pho84-/-* cells as determined by the phosphorylation state of ribosomal protein S6 (P-S6) [62] (Fig. 3) while null mutants of the other Pi transporters did not show reduced P-S6. We concluded that Pho84 specifically contributes to TORC1 activation among the Pi transporters. TORC1 contributes to managing oxidative stress responses in *C. albicans,* and mutants in *PHO84* are known to be hypersensitive to oxidative stress [25, 26, 59]. Among single null mutants in each of the Pi transporters, only the *pho84*-/- mutant showed hypersensitivity to the superoxide inducer plumbagin (S2 Fig.).

**Fig. 3.**
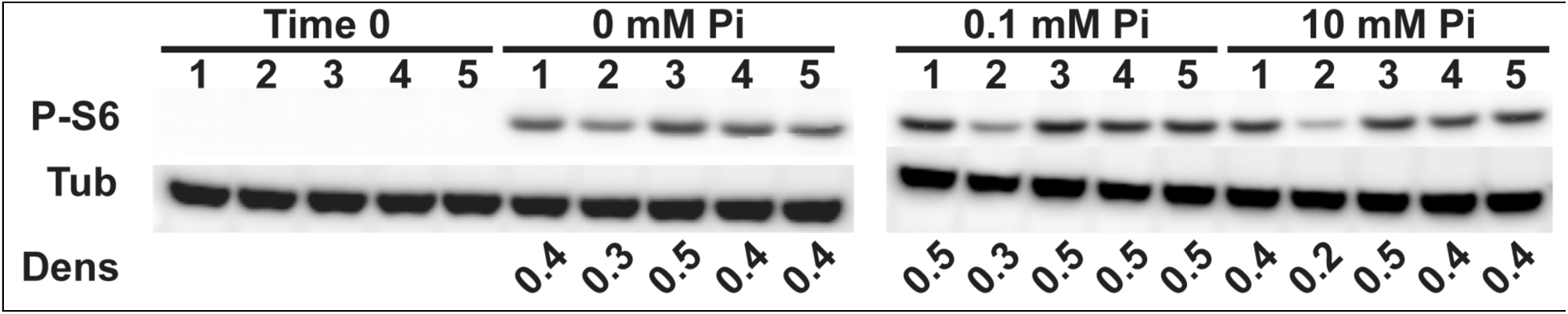
Pho84 was required for TORC1 activation. Cells were grown in YNB with indicated Pi concentrations for 90 min. Western blots were probed against phosphorylated Rps6 (P-S6) for monitoring TORC1 activity, and tubulin (Tub) as loading control. Dens: ratio between P-S6 and tubulin signals by densitometry. Representative of 3 biological replicates. Strains are 1: WT (JKC915); 2: *pho84-/-* (JKC1450); 3: *pho87-/-* (JKC2581); 4: *pho89-/-* (JKC2585) and 5: *fgr2-/-* (JKC2667)

### Triple and quadruple mutants in predicted Pi transporters support a major role for Pho84

To examine the Pi uptake characteristics of each transporter, we constructed triple mutants that retained one predicted Pi transporter, Pho84, Pho89, Pho87 or Fgr2. Triple mutants retaining a single Pi transporter are abbreviated as the name of the sole remaining transporter followed by a capital A for “alone among predicted Pi transporters” (e.g., Pho84-A is *pho89-/- pho87-/- fgr2-/-*). Among these triple mutants, only Pho84-A grew under all tested conditions (Fig. 1). Under Pi limiting conditions, Pho89-A cells showed significant growth only at neutral pH. These results support a role for Pho89 as a high-affinity transporter with a more alkaline optimum as in *S. cerevisiae* (Fig. 1). Pho87-A and Fgr2-A cells grew only in high Pi (7.3 mM) at acidic pH (pH 5 and pH 3) (Fig. 1), suggesting that Pho87 and Fgr2 are low-affinity transporters with an acidic optimum.

We engineered two quadruple Pi transporter mutants (Q-, *pho84-/- pho89-/- pho87-/- fgr2-/-*). These strains grew on rich complex medium, YPD, that contains organic phosphate compounds. They were then tested for growth on Pi as the sole phosphorus source at a range of concentrations. Q- cells were able to grow on high (7.3 mM) Pi at acidic pH, while on moderate (0.5 mM) Pi their growth was barely detectable (Fig. 1). They did not grow at pH 7 or at a low Pi concentration (0.05 mM, Fig. 1). These findings show that a residual Pi transport capacity exists in cells lacking the 4 identified Pi transporters that is active at high Pi concentrations and at an acidic pH.

### *PHO84* supported growth under the broadest range of conditions

We defined the pH range at which each triple mutant retaining a single predicted Pi transporter was able to grow in SC medium with low or high Pi. Growth was assayed by optical density and the area under each growth curve (AUC) was depicted as a histogram (Fig. 4 and S3 Fig.). Substantial growth was defined as growth ≥ AUC 5. Overall, we found that growth of Pho84-A (*pho89-/- pho87-/- fgr2-/-*) cells resembled that of WT (Fig. 4), with growth optima between pH 2 and pH 7 in both high and low Pi conditions.

**Fig. 4.**
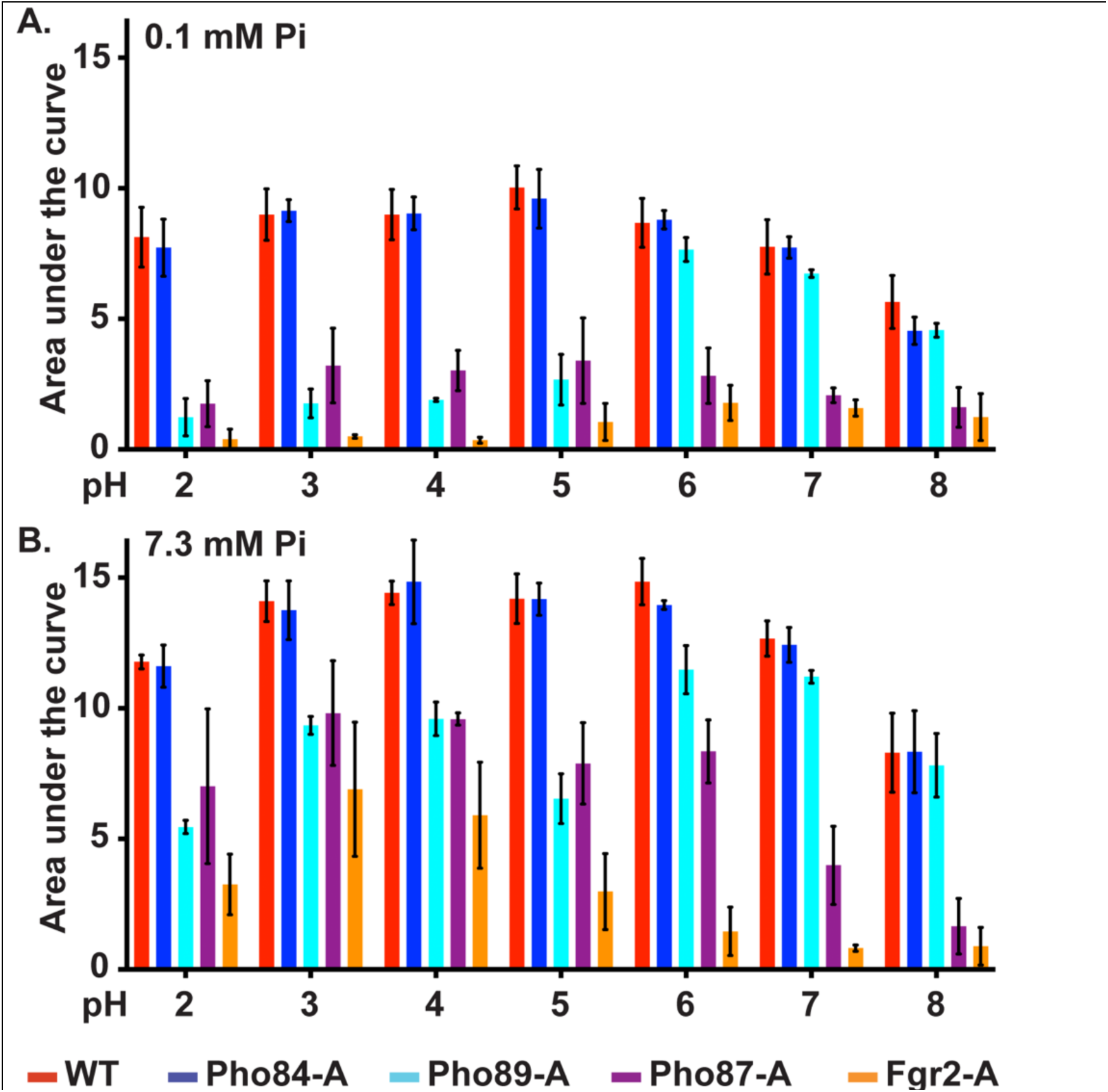
Pi transporters differed for their optimal pH and Pi concentration range while Pho84 was most active overall. Cells expressing the indicated Pi transporter alone among the 4 Pi transporters were inoculated to an OD_600_ of 0.2 into SC medium buffered to the indicated pH with 100 mM MES and grown in a plate reader at 30° C for 20 h. OD_600_ was measured every 15 min and the area under the growth curve was calculated in Graphpad Prism; means of 3 biological replicates are depicted in histograms. See S3 Fig. for the represented growth curves. Error bars represent SD of 3 biological replicates. **A.** SC containing 0.1 mM KH_2_PO_4_. **B.** SC containing 7.3 mM KH_2_PO_4_. Strains are WT (JKC915); Pho84-A: *pho87-/- pho89-/- fgr2-/- PHO84+/+* (JKC2788); Pho87-A: *pho84-/- pho89-/- fgr2-/- PHO87+/+* (JKC2777); Pho89-A: *pho84-/- pho87-/- fgr2-/- PHO89+/+* (JKC2783); Fgr2-A: *pho84-/- pho87-/- pho89-/- FGR2+/+* (JKC2758).

The role of Pho89 in growth was dependent on pH and Pi availability (Fig. 4). In low Pi, Pho89-A cells grew equivalently to WT at pH 6 and above. In high Pi, Pho89 supported intermediate levels of growth at more acidic pH and growth equivalent to WT at pH 6 and above. Pho89 could hence be described as a putative high-affinity Pi transporter in neutral and alkaline conditions and a low-affinity transporter in acidic conditions. *C. albicans*, like *S. cerevisiae*, therefore has two high-affinity Pi transporters, Pho84 and Pho89, with the former having a broad pH activity range including in alkaline conditions, and the latter active at pH ≥6.

Pho87 and Fgr2 were unable to support substantial growth in low ambient Pi and therefore are low-affinity Pi transporters. Both supported growth only at acidic pH in high Pi. Pho87-A cells grew to an AUC ≥5 only between pH 2 and 6, and even at their optimal conditions supported only ∼70% of the WT growth (Fig. 4). Fgr2-A cells showed the weakest growth with similar optima to Pho87-A cells, growing to an AUC ≥5 only at pH 3 and 4 (Fig. 4). *C. albicans* therefore has two low-affinity Pi transporters, Pho87 and Fgr2, with the latter, a Pho84 homolog, showing a narrow, acidic pH optimum.

Hyphal formation of triple Pi transporter mutants largely reflected the growth-sustaining properties of the transporters. Q-mutants lacking all four Pi transporters failed to form hyphae under any conditions tested (Fig 2). In contrast, Pho84-A cells encoding only Pho84 formed robust hyphae across all conditions (Fig. 2), consistent with the strong hyphal defect of *pho84-/-* mutants. Pho89-A cells had severe hyphal growth defects resembling those of *pho84-/-* mutants. Filamentation of the Pho87-A and Fgr2-A cells resembled the Q-mutants (Fig. 2). These mutants did not grow sufficiently to form hyphae on RPMI buffered to pH 7 (Fig. 2C). Collectively, these findings are consistent with a concept that hyphal growth requires Pi uptake.

### Pho84 showed the most active Pi uptake under all tested pH conditions

In order to quantify the Pi transport capacity of each transporter, we performed Pi uptake experiments with the triple mutants that each retained a single predicted transporter, by measuring Pi concentrations remaining in culture medium over a time course. WT and Pho84-A cells removed Pi from the medium rapidly and with almost identical kinetics (Fig. 5A,B). Pho89-A cells efficiently transported Pi at a narrow range of pH 6-8, but their uptake dramatically slowed at pH 5 and below (Fig. 5C,D). These results support a predominant role of Pho84 as the major Pi importer in *C. albicans,* while Pho89 makes a substantial contribution around neutral pH.

**Fig. 5.**
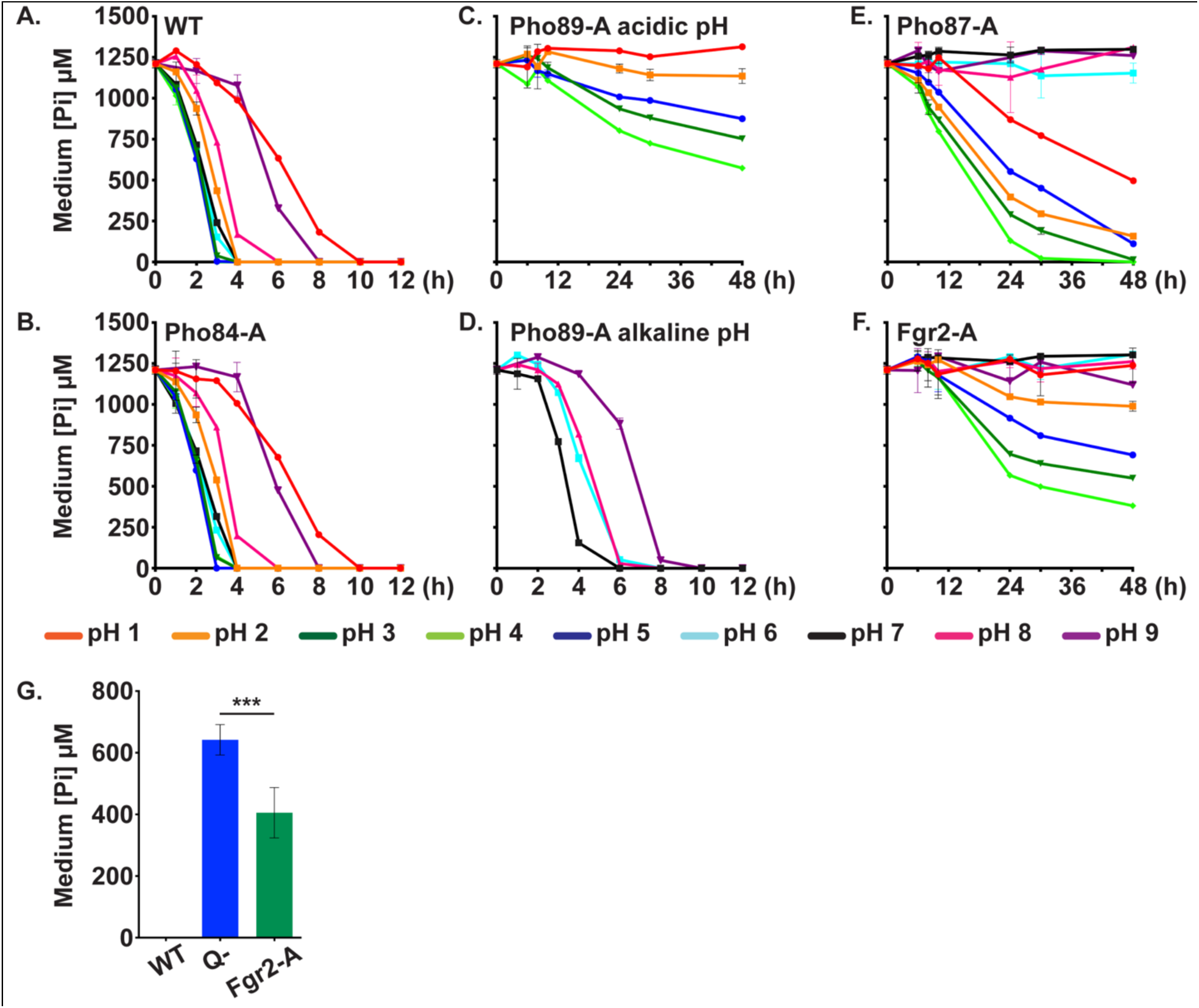
Pi uptake of cells expressing single Pi transporters reflected their growth optima. Cells with indicated genotypes were inoculated into SC without Pi (buffered to pH 1-9) at OD_600_ 2. After 30 minutes, KH_2_PO_4_ was added to a final concentration of 1 mM, and the extracellular concentration of phosphate was measured in 2 technical replicates at indicated time points. Error bars SD. Representative of three biological replicates. **A.** WT (JKC915). **B.** Pho84-A: *pho87-/-pho89-/- fgr2-/- PHO84+/+* (JKC2788). **C.** Pho89-A in pH 1-5; **D.** Pho89-A in pH 6-9: *pho84-/-pho87-/- fgr2-/- PHO89+/+* (JKC2783). **E.** Pho87-A: *pho84-/-pho89-/- fgr2-/- PHO87+/+* (JKC2777). **F.** Fgr2-A: *pho84-/- pho87-/- pho89-/- FGR2+/+* (JKC2758). **G.** Q- cells (*pho84-/- pho87-/- pho89-/- fgr2-/-;* JKC2830) took up significantly less Pi at pH 4 than Fgr2-A cells at 30 h of incubation. Histograms depict average and SD of 3 biological replicates, *p*=0.0001 (two- tailed t-test).

Low-affinity Pi transporters Pho87 and Fgr2 showed slow Pi uptake under these conditions. Uptake of Pi by Pho87-A cells was sluggish at pH 2-5 and almost undetectable at pH 6-9 (Fig. 5E). Despite known strong induction of *FGR2* in low Pi conditions by Pho4, the transcriptional regulator of the PHO regulon [61], Pi uptake by Fgr2-A cells was weak across the pH levels tested and these cells were unable to fully deplete Pi from the medium at their pH 4 optimum (Fig. 5F). Both Pho87-A and Fgr2-A cells removed Pi from the medium most rapidly at pH 4. Still, Pi uptake by Fgr2-A cells was significantly higher than that of Q-cells at 30 hours (*p*=0.0001 by two-tailed t-test, Fig. 5G). These data support a role for Pho87 and Fgr2 as Pi importers albeit with poor kinetics.

To identify any growth defects unrelated to Pi limitation in mutants containing a single Pi transporter, we compared growth of triple mutants to WT in liquid YPD and Pi replete SC media. YPD contains organic as well as inorganic phosphate sources, and SC contains high Pi concentrations (7.3 mM) and has an acidic pH of 4-5 that favors activity of most Pi transporters. No triple mutants exhibited a growth defect in YPD medium. In SC medium, Pho84-A grew as well as WT while Pho87-A, Pho89-A and Fgr2-A cells grew more slowly (S4 Fig.), consistent with our previous results (Fig. 1 for SC medium with 7.3 mM Pi at pH 5). These findings indicate that growth defects in these mutants correspond to a lack of Pi and not a nonspecific fitness loss.

### Glycerophosphocholine transporters provided a minor Pi import function

To test whether *C. albicans* expresses additional Pi transporters that were not detected by our homology searches, we next examined Q-cells for their ability to import Pi. At 30 hours, Q-mutants took up ∼40% of the Pi at pH 4 with efficiency being further reduced at pH 2, 5, 6 and 7 (Fig. 6A). These results demonstrated that another low-capacity Pi transporting activity existed at low pH and high Pi concentrations.

**Fig. 6.**
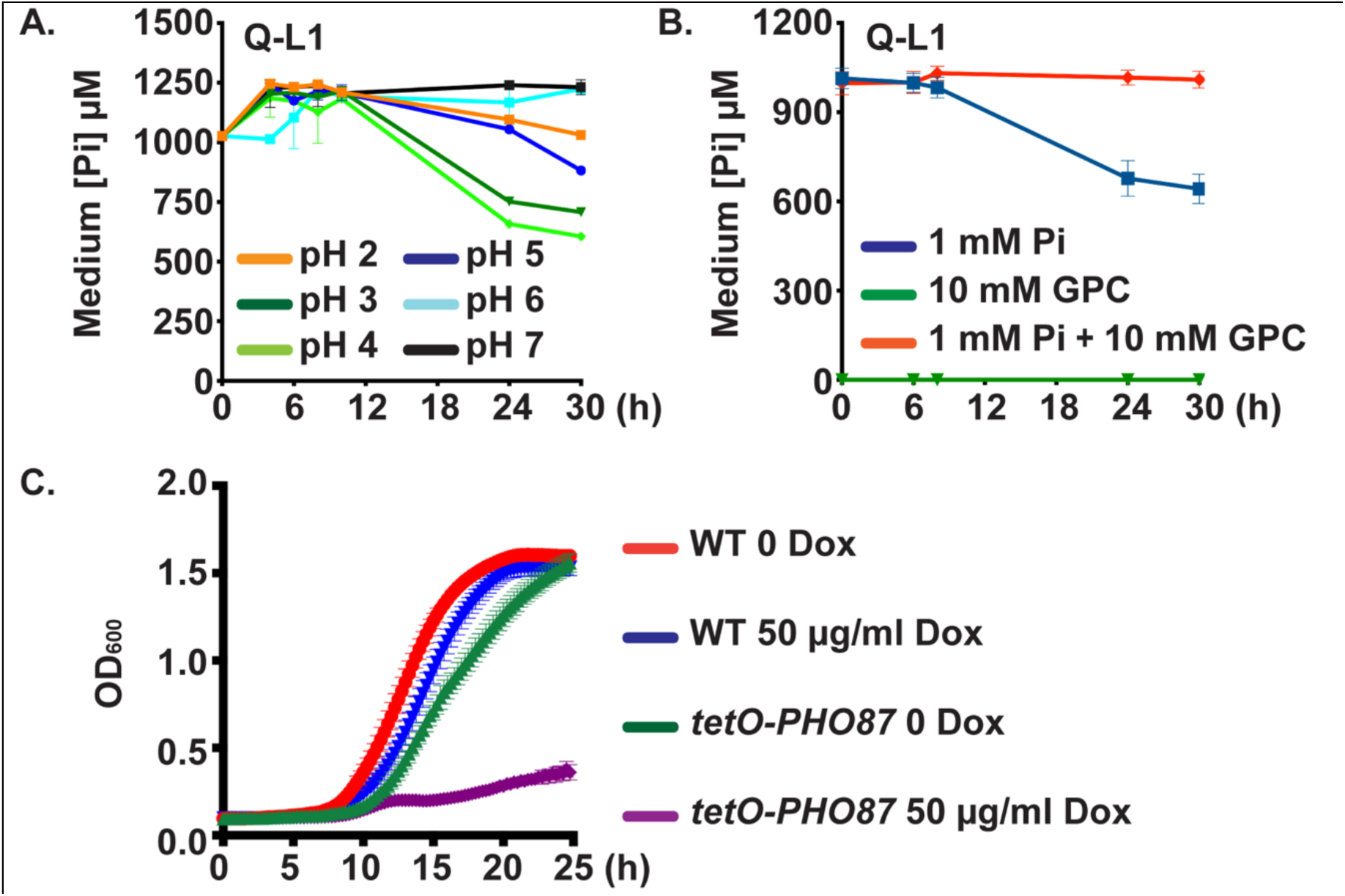
Cells lacking 4 Pi transporters showed residual Pi uptake ability that was outcompeted by glycerophosphocholine. **A.** Pi uptake experiments performed as in Fig. 5 showed that in Q-cells (*pho84-/- pho87-/- pho89-/- fgr2-/-*, JKC2830) residual Pi uptake occurred and was most efficient at pH 4. **B.** Tenfold excess glycerophosphocholine (GPC) inhibited Pi uptake in Q- cells at pH 4. As in Fig. 5, Q- cells (JKC2830) were inoculated into SC 0 Pi with 10 mM GPC; after 30 minutes, KH_2_PO_4_ was added to a final concentration of 1 mM; Pi concentration in the medium was measured with 3 technical replicates at each time point. Graph shows mean of 3 biological replicates. Error bars SD. **C.** Cells in which *PHO87* is expressed from repressible *tetO* while the other 3 Pi transporters and *GIT2-4* are deleted (*tetO- PHO87/pho87 pho84-/- pho89-/- fgr2-/- git2-4-/-,* JKC2969), grew well in the absence of doxycycline but grew minimally during *tetO* repression in 50 µg/ml doxycycline. WT (JKC915) and *tetO-PHO87* (JKC2969) were starved for Pi in SC 0 Pi in the presence of 50 µg/ml doxycycline for 48 h. The medium and doxycyline were replaced every 24 h. Cells were then inoculated at OD 0.01 into SC medium (7.3 mM Pi), buffered to pH 3 with 100 mM MES, without and with 50 µg/ml Doxycycline. OD_600_ was recorded every 15 min. Error bars SD of 3 technical replicates. Representative of 3 biological replicates.

*C. albicans GIT3* and *GIT4* are distant *PHO84* homologs whose products import glycerophosphocholine (GPC), a phospholipid degradation product that can serve as an organic source of phosphorus [63]. Once in the cytosol, GPC is metabolized to glycerol, choline and phosphate under Pi limiting conditions [63]. The GPC transporters Git3 and Git4 share 23% amino acid identity with Pho84 (41% and 39% similarity, respectively, S1 Fig.). To test whether Git3 and Git4 might contribute to the residual Pi transport in Q-cells, we competed Pi uptake by Git3 and Git4 with excess GPC in the medium. Addition of a 10-fold excess of GPC completely eliminated Pi uptake by Q-cells (Fig. 6B). These findings support a role for Git3/4 in Pi import at high ambient Pi and acidic pH.

We asked whether *C. albicans* has another modality to import Pi from its surroundings, in addition to the 2 high-affinity Pi transporters Pho84 and Pho89, the 2 low-affinity Pi transporters Pho87 and Fgr2, and the GPC transporters Git3 and Git4. We engineered a septuple mutant strain “*tetO-PHO87”* that lacked three Pi transporter homologs (*pho84-/- pho89-/- fgr2-/-*), GPC transporters that occupy adjacent loci on chromosome 5 (*git2-/- git3-/- git4-/-*), and had a single tetracycline-repressible allele of *PHO87* (*pho87-/tetO-PHO87)*. Mutants were maintained without doxycycline to retain maximal expression of *PHO87*. A role for Git2 in Pi import is currently not known; *GIT2* was deleted alongside the other two transporters in their initial characterization and it was included in this construct to permit comparisons with mutants described in Bishop et al. [63].

We reasoned that if a Pi-transporting activity remained in these cells, they would grow in Pi as their only source of phosphorus, both in the absence and presence of doxycycline, i.e., during induction and repression of *PHO87*. On the other hand, if we had mutated all transporters capable of importing sufficient Pi to sustain growth, doxycycline exposure in SC media, devoid of organic phosphate sources, would repress growth once internal Pi stores were depleted. We observed the latter result for the *tetO-PHO87* septuple mutants (*pho87-/tetO-PHO87 pho84-/- pho89-/- fgr2-/- git2-/- git3-/- git4-/-*) (Fig. 6C). In contrast at pH 3, an optimal pH for Pho87 activity, these mutants grew robustly in media without doxycycline, YPD or normal SC (which contains high Pi, 7.3 mM) though they had a slight growth defect compared to WT (Fig. 6C). Thus, the reduced growth of these mutants is largely attributable to Pi starvation and not a consequence of a nonspecific fitness loss (S5A Fig.). We concluded that minimal Pi import activity remains when *PHO84*, *PHO89*, *PHO87*, *FGR2* and *GIT2-4* have been genetically eliminated.

### In vitro evolution during Pi starvation restored growth in two distinct quadruple Pi transporter mutant lineages

We reasoned that in vitro evolution of *C. albicans* Q-mutants lacking all four Pi transporter homologs (*pho84-/- pho89-/- pho87-/- fgr2-/-*) could reveal adaptive mechanisms that facilitate growth during Pi scarcity. Our previous work showed that *pho84* null mutants’ cell wall contained less phosphomannan and had a thinner outer layer [26], suggesting that modifying cell wall structures while reserving scarce Pi for essential processes like nucleic acid biosynthesis could sustain Pi-starved cells. We propagated two lineages of the Q- mutants through 30 serial passages every other day in liquid SC medium with a moderate concentration of 0.4 mM Pi. To reduce genetic bottlenecks, a large number of cells (∼3.5×10^6^ cells in 10 μl of saturated culture) were reinoculated into 10 ml of fresh medium. The population at each passage was saved for DNA extraction and as a glycerol stock. During passaging, the growth rates of both Q- lineages increased substantially and in distinct increments. Growth rates of the Q- lineages ultimately plateaued before the end of the experiment. However, fitness recovery in the two lineages during passaging differed significantly. Lineage 1 (Q- L1), evolving from JKC2830, recovered fitness more rapidly and attained a growth rate similar to the WT by passage 24 (Fig. 7A). In contrast, growth rates of populations derived from lineage 2 (Q- L2, evolving from JKC2860) remained lower than WT throughout the experiment, plateauing at passage 20 (Fig. 7A). Fitness loss of Q- strains and fitness recovery of their descendant passaged populations was specific to Pi scarcity. In YPD, both Q- strains and their 30^th^ passage descendant populations grew robustly (S5B Fig.). These findings indicated that *C. albicans* cells lacking Pi transporters could recover fitness during prolonged Pi deprivation, and that this adaption did not lead to fitness loss when the selective pressure was relieved.

**Fig. 7.**
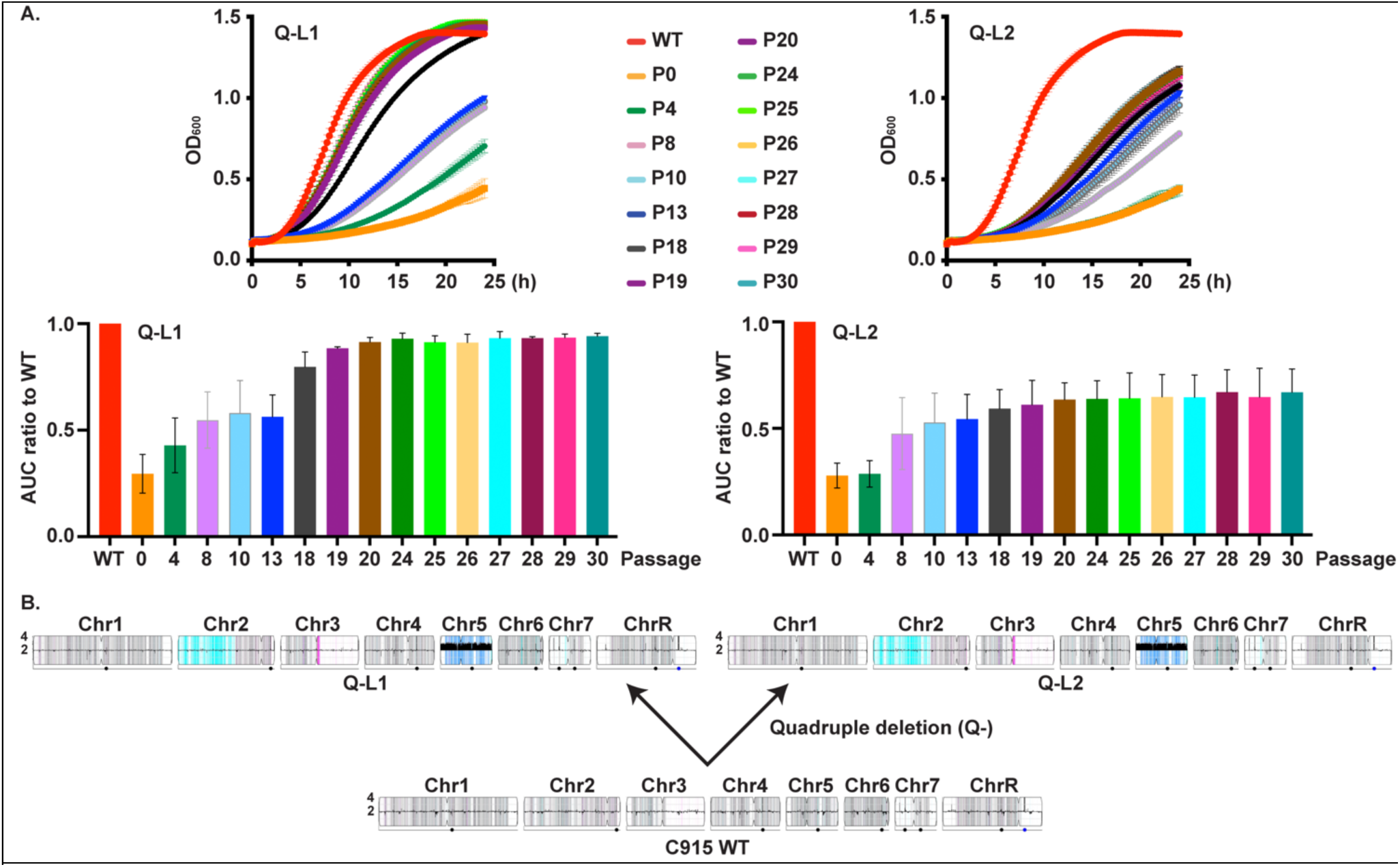
Whole genome sequencing revealed aneuploidies in Q-cells, and evolution of 2 Q-lineages during Pi scarcity proceeded along distinct trajectories. **A.** Cells from selected passages of populations evolving under Pi starvation from Q-isolates JKC2830 and JKC2860 were grown in SC 0.4 mM Pi. Representative growth curves and corresponding area under the curve (AUC) shown for selected passages. P0 denotes the Q-isolate before passaging; all experiments after P0 were performed with populations, not with clones derived from single colonies. Growth curves are representative of 3 biological replicates, except for passages 4 and 8, which are representative of 2 biological replicates. Error bars SD of 3 technical replicates. **B**. YMAP [74] depictions of WGS results of 2 distinct Q-isolates (*pho84-/- pho87-/- pho89-/- fgr2-/-*, JKC2830, Q- L1 and JKC2860, Q- L2) showing Chr5 trisomy and loss of heterozygosity of Chr2 and Chr3.

### Acquisition and loss of aneuploidies accompanied adaptation of evolving Q- lineages to Pi scarcity

To uncover the underlying genomic changes associated with improved growth of evolving Q- strains, we performed whole genome sequencing of the WT ancestral to the Pi transporter mutants (JKC915), as well as cell populations of selected passages from both lineages, Q- L1 and Q- L2. Populations that bounded significant fitness gains were chosen for sequencing, generating genomic snapshots of the evolved population for passages 4, 13, 18, 19, and 30 of Q- L1, and passages 5, 7, 19, 24, and 30 for Q- L2.

Major chromosomal rearrangements occurred during construction of the Q- mutants and during their passaging. The WT strain used to construct the Q- mutant lineages was diploid with no evidence of segmental copy number variations or large loss of heterozygosity (LOH) tracts. The genomes of both Q- strains were trisomic for chromosome 5 (Chr5, Fig. 7B), which includes *GIT2-4*. This aneuploidy was accompanied by LOH tracts on the left arm of Chr2 (Chr2L) and Chr3R, neither of which include loci disrupted in the Q- mutant. LOH of Chr2 resulted in homozygosis of the A allele for >50% of the chromosome, and LOH of Chr3 produced a short tract of homozygosity for B alleles (Fig. 7B).

Both evolving lineages acquired similar large-scale genomic changes during adaptation to Pi scarcity. Each lineage independently acquired a Chr2 trisomy, first seen in passage 13 and passage 19 for Q- L1 and Q- L2, respectively (Fig. 8). The Chr2 trisomy was subsequently largely or partially lost in Q- L1 and Q- L2, respectively, possibly as populations accumulated fitness-enhancing small-scale mutations. The Chr5 trisomy was also lost during passaging of Q- L1, likely due to fitness defects associated with aneuploidy (Fig. 8).

**Fig. 8.**
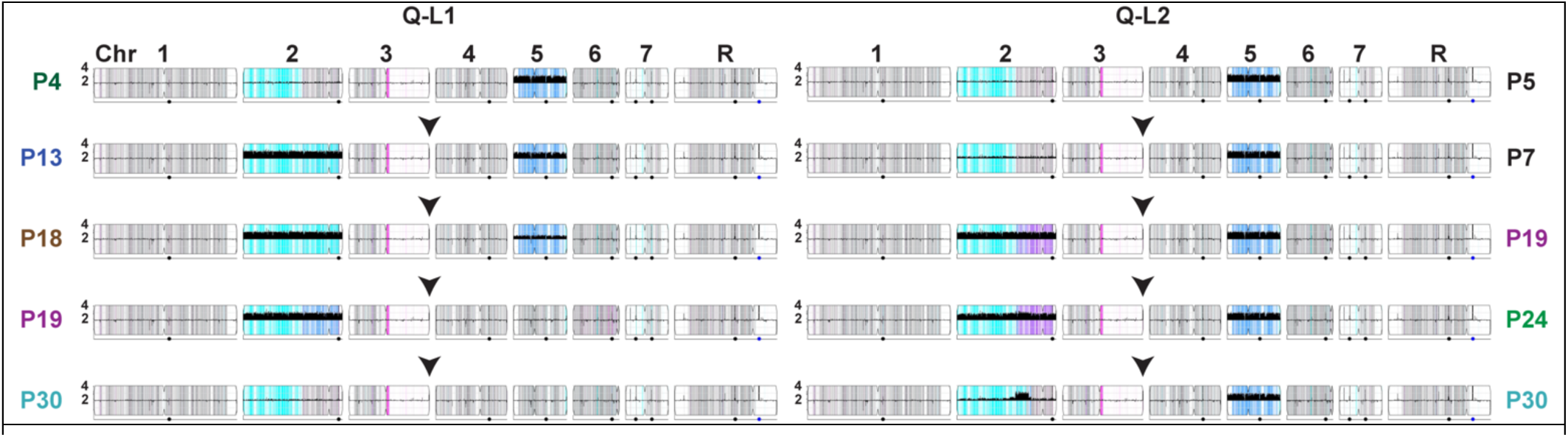
Whole genome sequencing of selected passages of 2 evolving Q- derived populations showed distinct trajectories of acquisition and resolution of aneuploidies and loss of heterozygosity. YMAP [74] depictions of WGS results of populations evolving from JKC2830, Q- L1 and JKC2860, Q- L2 (both *pho84-/- pho87-/- pho89-/- fgr2-/-*).

Persistence of Q- L2’s segmental aneuploidy on the left arm of Chr2 (Fig. 8) by the 30^th^ passage suggests potential adaptive contributions of the encoded loci. This amplified region of Chr2 ranged from nt 1,316,396 to 1,617,025 and encompassed 147 open reading frames. Manual annotation of these genes (SI Table 1) identified *GIT1,* which encodes a glycerophosphoinositol permease [64] with 22% amino acid sequence identity to Pho84. There is experimental evidence against a role of Git1 in Pi transport in *C. albicans* [64] but its role in phosphorus homeostasis might be indirect, i.e. by facilitating glycerophosphoinositol uptake as Pi starvation induces plasma membrane remodeling [65].

**Table 1.**
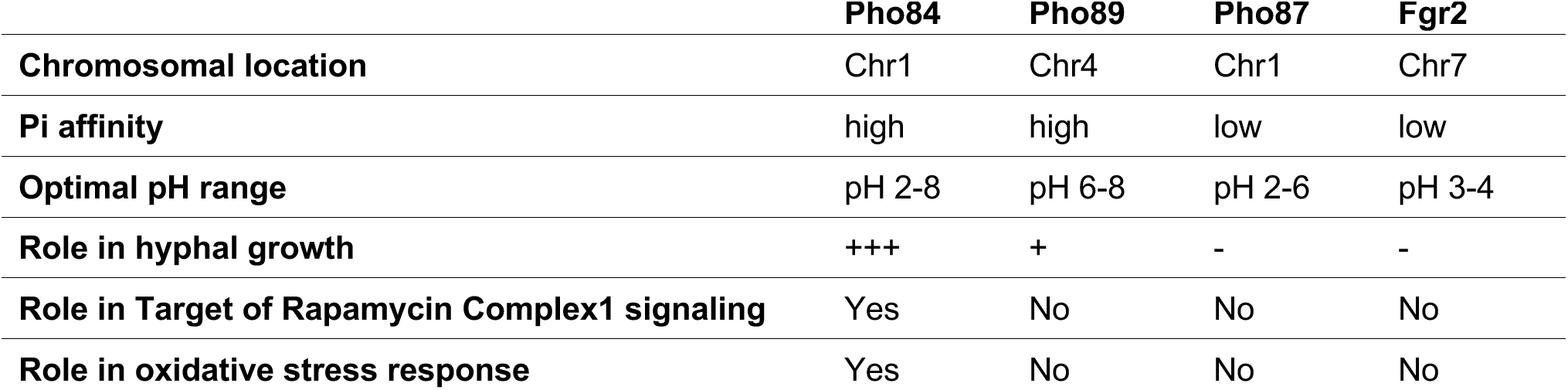
Pi transporter characteristics.

### Specific stress phenotypes of Q- population passages did not consistently correspond to their fitness in Pi scarcity

As we previously found hypersensitivity of *pho84* null mutants to rapamycin as well as oxidative-, cell wall- and membrane stress, induced through plumbagin, micafungin and amphotericin exposure respectively [24, 26], we investigated these responses in the Q- mutants and their evolved lineages. WT, both Q- strains and their passage 30 descendant populations were grown in liquid medium; these strains and in addition, populations from intermediate passages from the evolution experiment were spotted as serial dilutions onto solid medium in the absence or presence of these stressors. Q- mutants were hypersensitive to membrane stress induced by amphotericin and SDS and to cell wall stress induced by micafungin; they showed rapamycin sensitivity corresponding to their growth defects in vehicle (Fig. 9).

**Fig. 9.**
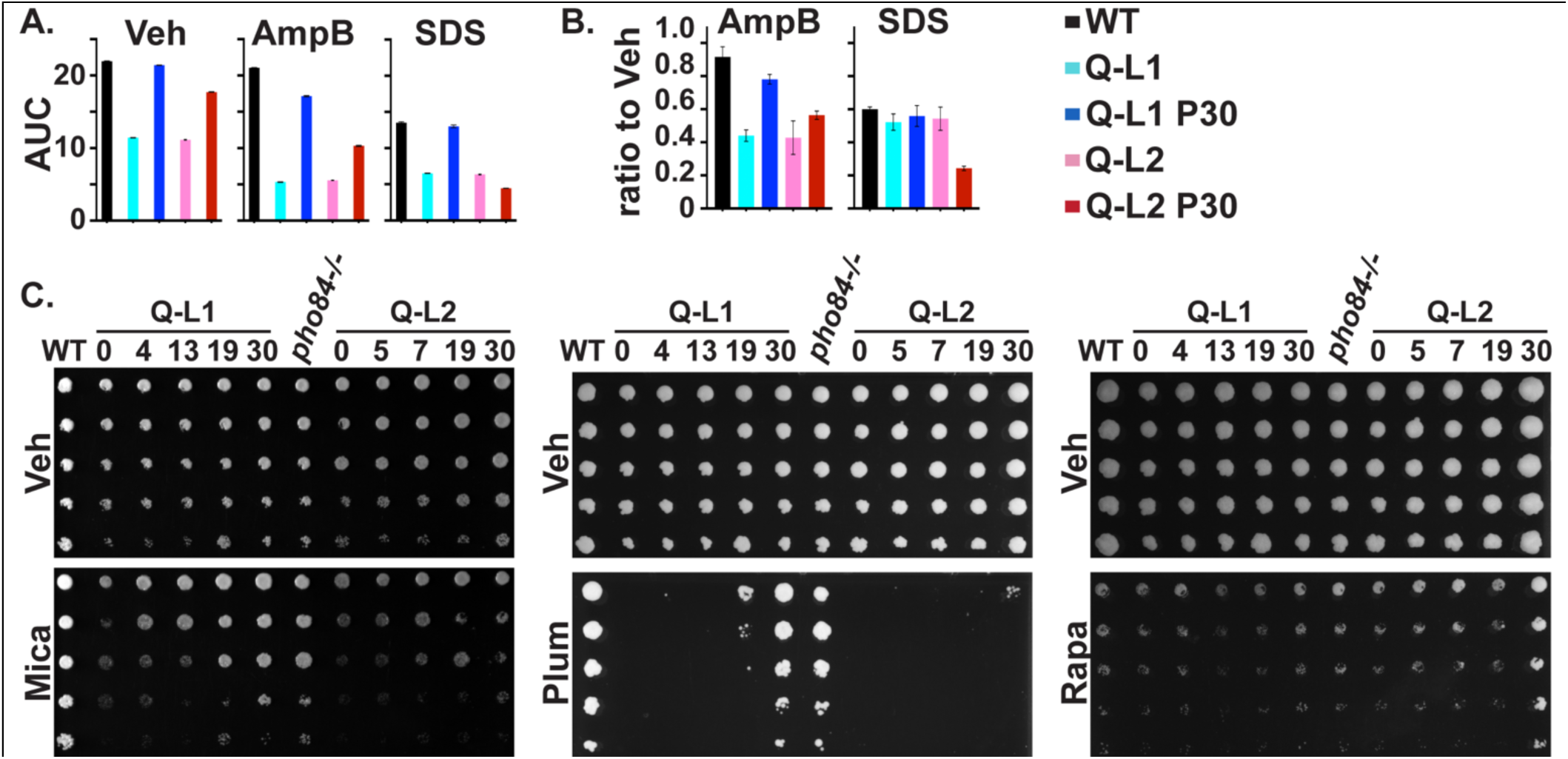
Two evolving lineages of Q- cells showed distinct stressor endurance. **A**. Growth area under the curve (AUC) of cells grown in SC 1 mM Pi containing Vehicle (Veh, DMSO), 0.3 µg/ml Amphotericin B (AmpB) or 0.005% SDS. AmpB representative of 2 and SDS of 3 biological replicates; error bar SD of 3 technical replicates. **B**. Amphotericin B (AmpB) and SDS growth area under the curve (AUC) from panel A normalized to each strain’s vehicle (Veh) control. WT (JKC915); Q- L1 (JKC2830); Q- L1 P30 (JKC2830 passage 30); Q- L2 (JKC2860); Q- L2 P30 (JKC2860 passage 30). AmpB average of 2 and SDS of 3 biological replicates; error bar SD of biological replicates. **C**. Threefold dilutions of cells of indicated genotypes, starting at OD_600_ 0.5, were spotted (top to bottom) onto SC medium containing Vehicle (Veh, H_2_O) or 10 ng/ml micafungin (Mica), 15 µM plumbagin (Plum), 50 ng/ml rapamycin (Rapa), and grown at 30°C for 1 d (Mica), 2 d (Plum), 4 d (Rapa), respectively. Strains are WT (JKC915); Q- L1 (JKC2830) passage 0, 4, 13, 19, 30 and Q- L2 (JKC2860) passages 0, 5, 7, 19, 30; *pho84-/-* (JKC1450). JKC2830 and JKC2860 genotypes are *pho84-/- pho87-/- pho89-/- fgr2-/-*.

In contrast, the responses of evolved, late-passage Q- L1 and Q- L2 populations were distinct for each stressor. Q- L1 but not Q- L2 populations regained growth rates in amphotericin almost to WT levels by passage 30 (Fig. 9A,B). The Q- L2 passage 30 population had increased sensitivity to SDS, compared with its ancestral Q- L2 strain (Fig. 9A,B) while its Q- L1 counterpart regained the ability to grow in the presence of SDS almost to the level of the WT (Fig. 9A,B). In the presence of micafungin, growth of selected passages reflected their fitness in Pi-limited medium (Fig. 9C). However, Q- L1 but not Q- L2 passaged populations regained growth in plumbagin while conversely, Q- L2 populations evolved frank resistance to rapamycin by passage 30 (Fig. 9C). Like the different fitness levels in Pi scarcity reached by the end of the experiment, the distinct stress phenotypes of Pi scarcity-adapted populations suggest their evolutionary trajectories had diverged.

## Discussion

In this work, we characterized 4 predicted *C. albicans* Pi transporters, Pho84, Pho89, Pho87 and Fgr2, identified by sequence homology, and determined their contributions to Pi acquisition. In brief, among the high-affinity Pi transporters, Pho84 was the most important for growth, filamentation, stress responses, and induction of TORC1 signaling and had the broadest pH range of Pi uptake capacity, while Pho89 was specialized for uptake in neutral and alkaline pH (Table 1). Among the low-affinity Pi transporters, both of which were only active at acidic pH, Pho87 was more efficient and had a broader pH range while Fgr2 functioned only between pH 3 and 5 (Figs. 4, 5). In contrast to *S. cerevisiae*, *C. albicans* low-affinity Pi transporters are not paralogs; rather, the less efficient one, Fgr2, is a distant Pho84 homolog. The minor role for Fgr2 in *C. albicans* Pi import stands in contrast to *Cryptococcus neoformans* where both *PHO84* homologs (*PHO84* and *PHO840*) make significant contributions to Pi transport [66]. In the absence of all specific Pi transporters, glycerophosphocholine transporters were able to provide residual Pi import to sustain growth. Cells lacking all Pi transporters were able to regain fitness during sequential passaging in limited Pi, that plateaued at distinct levels for 2 populations in accordance with previously described declining adaptability [67, 68], while engendering distinct responses to some Pi- relevant stressors. During the evolution experiment, similar large-scale genomic changes were sequentially acquired and partially or completely lost in the 2 independently evolving populations. Retention of a segmental amplification within Chr2 in the less fit lineage suggests that genes in this trisomic region may contribute to fitness in limited Pi but the precise loci responsible remain undefined.

The severe growth defect of Q- cells that lack all 4 identified Pi transporters argues against the presence of other specific Pi transporters. Q- cells removed a small fraction of the Pi present in their medium. The glycerophosphocholine transporters Git3 and 4 provided minor transport activity that was most evident in the growth difference between a Q- mutant and the septuple mutant when grown in the presence of doxycycline to repress *PHO87* (Fig. 6C). The ability of GPC provision to completely outcompete measurable Pi uptake in Q- cells (Fig. 6B) also argues against other cell surface transporters beside Git3 and Git4 playing a measurable role in Pi uptake under our experimental conditions.

pH sensitivity is a key feature of each transporter. Like WT, Pho84-A, Pho87-A, and Fgr2-A cells grew optimally between pH 3 and 6. Only cells expressing *PHO89* alone showed a growth optimum between pH 6 and 8. The pH of the oral and pharyngeal mucosa as well as most of the gastrointestinal tract colonized by *C. albicans* is broadly neutral or alkaline, though mucosal microenvironments may be acidic due to bacterial metabolites. During acute invasive disease, *C. albicans* finds itself in mildly alkaline blood and tissue environments between pH 7.35 and 7.45. Pi acquisition systems of *C. albicans* are therefore not well adapted to host environments encountered during invasive disease. The reduced Pi transport activity at alkaline pH of the bloodstream might explain why the PHO regulon is induced during systemic disease, reflecting “alkaline pH-simulated nutrient deprivation” [69], despite the presence of abundant Pi and organic phosphate compounds like GPC. Pi import is also critical for proliferation of other human fungal pathogens [30, 70, 71] and unicellular parasites [72] each of which must contend with neutral to alkaline conditions in host deep organs.

Loss of *PHO84* but not of the other Pi transporters had a substantial effect on TORC1 activity (Fig. 3) and oxidative stress endurance (S2 Fig.). These experiments cannot distinguish between specific activities of Pho84 in these cellular functions, versus the predominant role of Pho84 in providing Pi to the cell. We found in another context that TORC1 activity depends on availability of Pi but not on the presence of Pho84, while endurance of peroxide stress may require an activity specifically of Pho84 [73]. *C. albicans PHO84* transcription is co-regulated by TORC1 in addition to Pho4 [24], but how these systems interact and modulate each others’ outputs remains unknown.

Hyphal growth defects mirrored the severity of Pi transport deficiency among the constructed mutants. Among the Pi transporters assayed in defined mutants, only Pho84-A cells produced robust hyphae on all 3 media examined (Fig. 2), while Pho87-A cells produced some hyphae on RPMI pH 5, and Pho89-A cells had sparse, short hyphae on RPMI pH 7 (Fig. 2). The role of Pi uptake in filamentation might be indirect, through activation of signaling systems like TORC1 required for hyphal morphogenesis [74]. Alternatively, given the larger surface area of hyphal cells compared to yeast cells, hyphal cells must consume larger amounts of phosphoric metabolites like nucleotide sugars required as building blocks for the cell wall. The inability to produce sufficient phosphorus-containing intermediates might inhibit hyphal morphogenesis to minimize cell wall surface area and preserve phosphorus for other vital functions.

To probe *C. albicans’* options for adaptation to Pi scarcity, we performed an in vitro evolution experiment with Q- strains in which we had deleted the known Pi transporters. These strains had acquired, at an unknown point during their construction, a trisomy of Chr5 where the genes encoding organic phosphate transporters Git3 and Git4 reside. *C. albicans* GPC transporters are upregulated by the transcription factor Pho4 during Pi starvation [61]. As Q- cells grew very poorly in medium with a moderate Pi content of 0.4 mM, increasing the gene dosage of *GIT3* and *GIT4* by retaining a third copy of Chr5 still did not restore significant Pi uptake, as shown in Fig. 5G. During the evolution experiment, a further large-scale genomic alteration appeared in both lineages: triploidy of Chr2 with simultaneous LOH reducing all 3 alleles to AAA in a long segment on the left arm of the chromosome (cyan-colored segment of Chr2 in both lineages, Fig. 8).

When abruptly exposed to significant stress, *C. albicans* is known to frequently resort to aneuploidy and LOH to provide a crude but rapid option to improve fitness specific to the particular stress [48–50, 75, 76], reviewed in [77]. Ploidy increases that enhance fitness under specific stress conditions can be achieved more rapidly than accumulation of advantageous (in the setting of the specific stress) point mutations because chromosome missegregation occurs once every 5 × 10^5^ cell divisions (in *S. cerevisia*e) [78] while substitution of any particular base pair is estimated to occur once every 1.2 × 10^10^ cell divisions in *C. albicans* [79] and every 1.67 × 10^10^ cell divisions in *S. cerevisiae* [80]. Populations under strong selection are therefore more likely to initially become enriched for aneuploid mutants than for cells containing constellations of advantageous point mutations [81]. Large-scale copy number variants like trisomies however incur fitness costs due to increased transcription and translation of a multitude of genes that result in excess protein production and protein complex stoichiometry imbalances [78, 82, 83]. These fitness costs predominate when the selective pressure of the stress relents [48, 83]. In addition to environmental changes that diminish stress intensity, small-scale genomic changes (like single nucleotide mutations and small insertions or deletions) that promote adaptation to the specific stressor can relieve selective pressure and favor loss of a trisomic chromosome. For example, a gain-of-function point mutation of a transcriptional regulator [48, 84], may alleviate a specific stressors’ selective pressure to the point that trisomies resolve back to diploid, as we observed in the Q- L1 but not the Q- L2 lineage.

Evolving quadruple Pi transporter mutants similarly gained trisomies in both independent populations, highlighting the strong selection experienced by these cells. Loss of trisomies during passaging may have been enabled by accumulating fitness-enhancing small-scale variants. The distinct mutations that permitted these adaptive solutions remain to be identified. Given their cell wall- and membrane stress response phenotypes, it is tempting to speculate that Pi-sparing modifications of membranes and the cell wall might increase fitness during Pi scarcity. The cyanobacterium *Prochlorococcus,* a dominant species in waters of the North Pacific Subtropical Gyre, has a competitive advantage in this Pi-scarce ecosystem due to its membrane composition largely of sulfo- and glycolipids, in which fatty acids are linked to sulfate/sugar- or sugar-based, instead of phosphate-based, polar head groups [85]; other phytoplankton use similar adaptations [4]. *C. albicans* also economizes on non-essential uses of Pi by remodeling its membrane systems: the gene encoding a homolog of diacylglyceryl-*N*,*N*,*N*-trimethylhomoserine synthase is strongly upregulated during Pi starvation in dependence on the PHO regulon [61] and like *Neurospora crassa* and *C. neoformans, C. albicans* can replace membrane phospholipids with betaine-headgroup lipids during Pi starvation [65, 86].

A caveat is that the distinct stress phenotypes of the two passage 30 populations could be incidental or integral to their Pi management strategies. Detailed genotype comparisons and further genetic and cell biologic analysis of distinct mutations in the two lineages will be required to test causal relationships between the phenotypes. For example, while distinct SDS stress phenotypes and distinct strategies to Pi scarcity adaptation could be unrelated, another possibility is that the overall genomic changes that occurred in the Q- L2 during Pi scarcity adaptation led to membrane changes that rendered it susceptible to detergent stress from SDS. In contrast, the adaptation trajectory of lineage Q- L1 may have required less changes to membrane composition so that its SDS endurance was restored as it adapted to growth in scarce Pi. Genetic, genomic and lipidomic analyses will be required to distinguish these possibilities.

These experiments have several limitations. As noted, there was a residual Pi transport activity in addition to the predicted Pi transporters present in triple mutant cells due to presence of GPC transporters Git3 and Git4; this residual activity was minor, though, and only detectable at high ambient Pi and acidic pH. Our experiments permit only semi-quantitative conclusions about the efficiency and optimum of each transporter, since we did not assay binding affinity and maximal transport velocity with ^32^P uptake measurements. Pi uptake experiments in early timepoints are likely not confounded by different growth rates of each triple mutant expressing a single Pi transporter “alone” because all experiments were performed at an OD_600_ of 2. However, inefficient Pi uptake may be artifactually amplified by slower growth of mutants at time points longer than 4-6 hours. Another potential confounder could be differential expression levels of the 4 transporters, which are likely to vary between pH and Pi concentration conditions; e.g. the alkali-responsive transcriptional regulator Rim101 induces transcription of *PHO89* [87], and protein levels of these 4 *C. albicans* transporters remain to be defined. Our in vitro evolution experiment was limited in that 2 transformants from a final strain-construction transformation step were evolved without technical replicates; nevertheless, experimental testing of candidate variants will be required in any case to ascribe fitness roles in Pi scarcity to mutations identified in both or in one of the phenotypically distinct lineages.

In summary, *C. albicans’* Pi acquisition system is suboptimal for the neutral to alkaline host environments it typically occupies during invasive disease. Differences in functional optima among transporters may provide backup mechanisms for Pi transport as *C. albicans* moves through different host niches. Redistribution of intracellular Pi among organelles and processes may sustain survival during “alkaline pH- simulated nutrient deprivation” [69], in ways that remain to be elucidated. At the same time, this redistribution might also render the fungus more sensitive to host-relevant stressors like membrane- and cell wall stress. Pi acquisition and regulation in humans differs fundamentally from that in fungi; given the crucial role of phosphorus in structural, metabolic and regulatory processes and in antifungal drug endurance, definition of these systems could reveal potential fungus-specific drug targets.

## Methods

### Culture conditions

Cells were grown in rich complex medium, YPD, defined media synthetic complete (SC) or yeast nitrogen base (YNB) with 2 % glucose as described in [25, 26]. To provide defined Pi concentrations, Yeast Nitrogen Base without Amino Acids, without Ammonium sulphate and without Phosphate, supplemented with KCl, was used (CYN6802, Formedium, Swaffham, Norfolk, England) and supplemented with the indicated concentrations of KH_2_PO_4_. Incubation temperatures were 30°C for liquid and solid media except for hyphal growth assays which were incubated at 37°C.

### Strain construction

*C. albicans* mutants were generated as in [88]; details of strain construction are provided in S2 Table (Strains used in this study), S3 Table (Plasmids used in this study) and S4 Table (Oligonucleotides used in this study). At least 2 and typically 3 independent heterozygous lineages were constructed for each set of deletion mutants. Only during construction of the Q- strain, 2 isolates were generated from a single *pho84-/- pho89-/- pho87-/- fgr2/FGR2* heterozygous parent. At each mutant construction step, we compared growth phenotypes of independent transformants in synthetic media; isolates with outlier phenotypes were eliminated from further use, since those presumably were due to unrelated mutations that arose during transformation. Among isogenic mutants with similar phenotypes, we performed initial phenotypic characterizations with 2 or more isolates to confirm their similar behaviors. The null mutants for each predicted transporter were used to examine the role of that transporter in filamentous growth, stress responses, susceptibility to antifungal agents and TORC1 signaling.

### Growth curves

Cells from glycerol stocks at -80°C were recovered on YPD agar medium for 2 days. Cells were scraped from the plate and washed twice with NaCl 0.9% and diluted to a final OD_600_ 0.01 in 150 µl medium in flat bottom 96-well plates. OD_600_ readings were obtained every 15 min in a Biotech Synergy 2 Multi-Mode Microplate Reader (Winooski, VT, USA). Standard deviations of three technical replicates, representing separate wells, were calculated, and graphed in Graphpad Prism Version 9.5.1 (528), and displayed as error bars. At least 3 biological replicates were obtained on different days unless stated otherwise. For some assays the area under the curve was calculated and graphed using the same software, and the average from ≥3 biological replicates per condition was graphed.

### Phosphate uptake

Phosphate uptake measurement experiments were performed as in [23] with some modifications. In brief, cells were incubated with a defined amount of Pi; the rate of Pi removal from the medium corresponds to the strains’ Pi transport capacity. Cells were recovered on YPD plates from glycerol stocks at -80°C for 2 days. SC medium without Pi, buffered at one-unit increments from pH 2 to 8 was inoculated with cells at an OD_600_ of 2. Cells were given 30 minutes to equilibrate, 1 mM Pi was then added, and the Pi remaining in the medium was measured every 1-5 h for a period of 6 to 30 hours, depending on each strain or condition. Samples were harvested at 20,000 rpm for 5 min in the cold room and two technical replicates per sample (300 µl) were collected from each time point. The total concentration of Pi in the supernatant was calculated according to Ames [89].

When the assay required pre-feeding the cells with GPC before the addition of Pi, 10 mM of GPC was added during the 30-minute incubation without Pi. Then Pi was added at a concentration of 1 mM.

### Growth of cell dilution spots on solid medium

Cells recovered from glycerol stock at -80°C were grown on YPD plates for ≥2 days. They were washed in 0.9 % NaCl and diluted in 5- or 3-fold steps from a starting OD_600_ of 0.5 in a microtiter plate, then pin transferred to agar medium and incubated for 48 h at 30°C.

### Hyphal morphogenesis assay

Cells recovered from glycerol stock at -80°C were grown on YPD plates for 1-2 days, washed and resuspended in 0.9% NaCl to an OD_600_ 0.1. Variation between spots and spot density effects were minimized by spotting 3 µl cell suspensions at 6 equidistant points, using a template around the perimeter of an agar medium plate. Each agar plate contained a WT spot that served as a control to which the other strains on the plate must be compared. This method minimizes variation between colony filamentation within each genotype that occurs when colonies are streaked or plated at varying density and at uncontrolled distances from each other. By including a WT on each plate, we also control for the effects of different hydration states of the agar and slight variations in medium composition which cannot be excluded by other means. RPMI and Spider media were used; the latter is not buffered and has a slightly higher than neutral pH. RPMI medium pH 7 was buffered with 165 mM MOPS; and RPMI pH 5 was buffered with 100 mM MES. Plates were incubated at 37°C for the indicated durations. All panels shown represent at least 3 biological replicates.

### Western blot

Cell lysis and Western blotting were performed as described in [62]. Antibodies used are shown in S5 Table. For densitometry, ImageJ (imagej.net/welcome) software (opensource) was used to quantitate signals obtained from Azure biosystems c600.

### Population evolution

Two Q- mutants (*pho84-/- pho89-/- pho87-/- fgr2-/-*), distinct transformants (Q- L1 and Q- L2)for deletion of the second *FGR2* allele, derived from the same heterozygous strain (JKC2812 *pho84-/- pho89-/- pho87-/- fgr2-/FGR2*), were inoculated into 10 ml SC 0.4 mM Pi at an OD_600_ of 0.05. Cultures were incubated at 30°C at 200 rpm. From each culture, 10 µl were transferred into 10 ml of the same fresh medium every 48 hours, for a total of 30 passages. The culture from each passage was saved as a glycerol stock. Populations from each passage were used in growth curve experiments and were always used after direct revival from frozen stock without further passaging.

### Genomic DNA isolation and whole-genome sequencing

For DNA extraction from cells grown on YPD plates from cells saved at the end of each passage, the Zymo Quick DNA Fungal/Bacterial Miniprep kit was used according to the manufacturer’s instructions.Library preparation and sequencing was carried out by the Applied Microbiology Services Lab (AMSL) at The Ohio State University, using the Illumina Nextseq 2000 platform to generate 150 basepair paired-end reads. All samples were sequenced to a minimum depth of 175x. The reads were trimmed using trimmomatic 0.39 (with default parameters except slidingwindow:4:20, maxinfo:125:1, headcrop:20, and minlen:35) to remove adaptors and poor quality sequences [90]. Using bowtie2 v2.2.6 [91], the trimmed reads were aligned against the SC5314 reference genome (version A21-s02-m09-r10) obtained from the Candida Genome Database (www.candidagenome.org). The aligned SAM files were then converted to the BAM format using samtools v1.7 [92].

### Copy number analysis

To detect karyotypic changes, pileup data for each whole-genome sequenced strain was obtained using bbMAP v39.01 [93]. Average read pileup depth highlighted any whole-chromosome aneuploidies. Copy number variation, including aneuploidies, were further confirmed via visualization in YMAP [94].

## Data availability

The sequencing data are available at the Sequence Read Archive under BioProject Accession Number PRJNA1035923.

## Acknowledgements

We thank Yuping Li for helpful discussions. M.A.-Z. was funded in part by the Alfonso Martin Escudero Foundation (Madrid, Spain). J.R.K. was funded by Boston Children’s Hospital and by NIAID R21AI096054 and NIAID R21AI137716. M.Z.A. was funded by National Institutes of Health grant 1R01AI148788 and an NSF CAREER Award 2046863. A.M. was supported by a President’s Postdoctoral Scholars Program Award at The Ohio State University. J.P.-V. was funded by NIH R15 GM104876. The funders played no role in the study design, data collection and analysis, decision to publish, or preparation of the manuscript. The authors declare no competing interests, financial or non-financial.

---

**S1 Fig.**
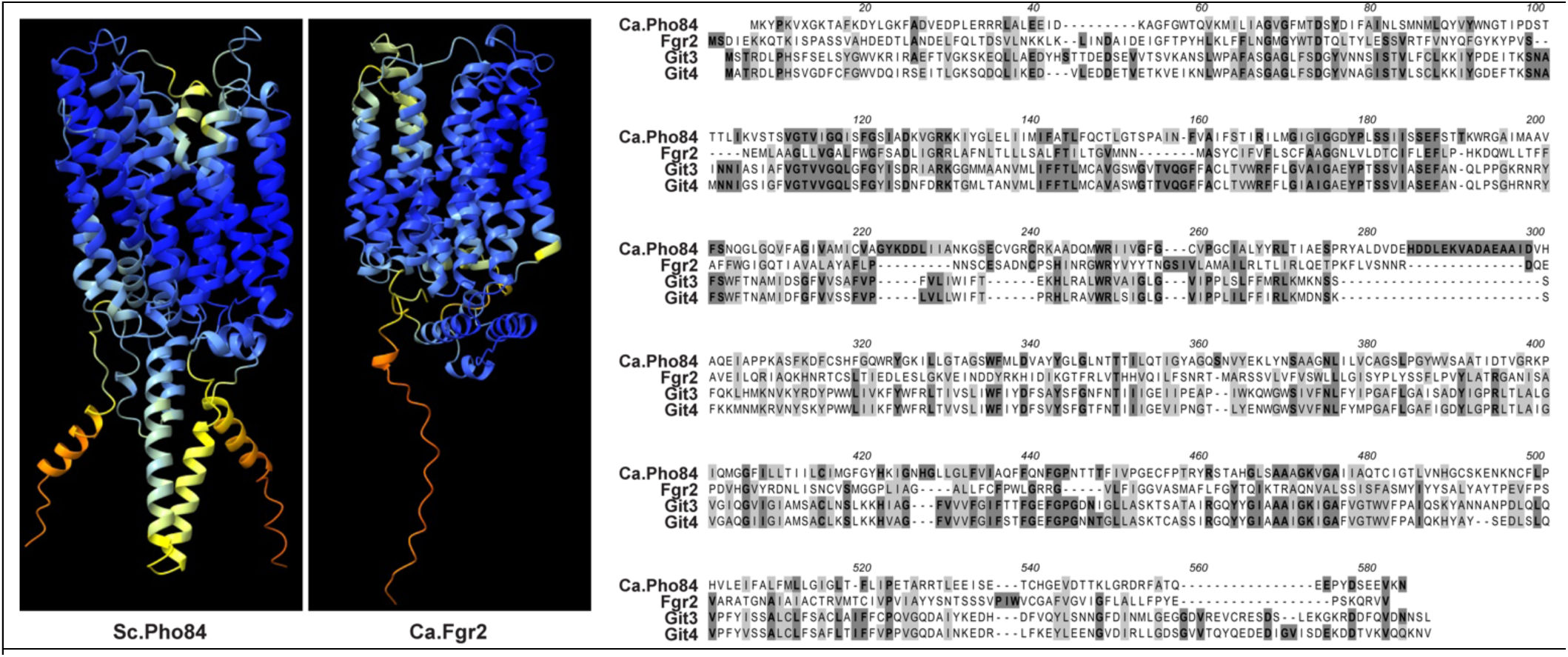
Structural comparisons of Pho84 and its homologs. AlphaFold structure prediction of *S. cerevisiae* Pho84 and *C. albicans* Fgr2 was obtained from the AlphaFold Protein Structure Database (https://alphafold.ebi.ac.uk/) and visualized in ChimeraX-1.4. The coloring of each model is based on a per-residue confidence score (pLDDT): dark blue – very high (pLDDT > 90), light blue – confident (90 > pLDDT > 70), yellow – low (70 > pLDDT > 50), orange – very low (pLDDT < 50). Inorganic phosphate transporters were aligned in MacVector; identical amino acid residues tinted with dark gray and chemically similar ones in light gray.

**S2 Fig.**
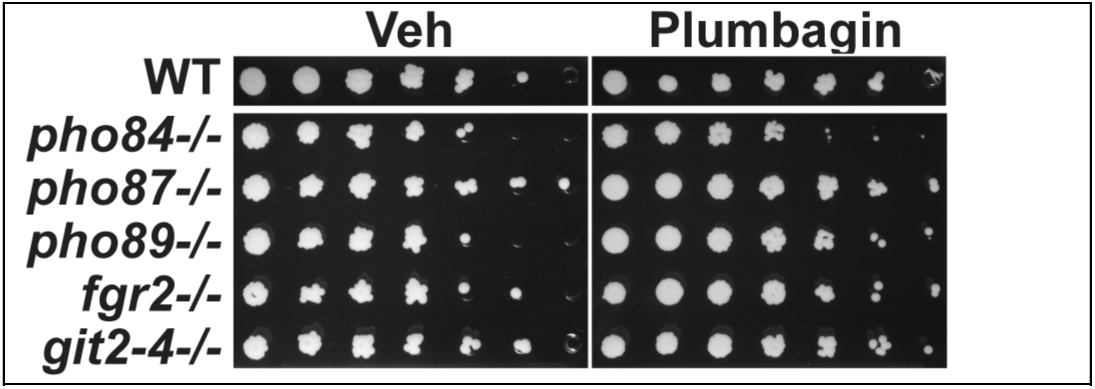
Among Pi transporters, only Pho84 was required for oxidative stress endurance. Cell suspensions of the indicated genotypes WT JKC915; *pho84-/-* (JKC1450); *pho87-/-* (JKC2581); *pho89-/-* (JKC2585); *fgr2-/-* (JKC2667) and *git2-4-/-* (JKC2963) were spotted in 3-fold dilution steps onto SC medium with DMSO (Veh) or plumbagin 15 µM. Plates were incubated for 2 days at 30° C. Representative of 3 biological replicates. All spots were on the same plate.

**S3 Fig.**
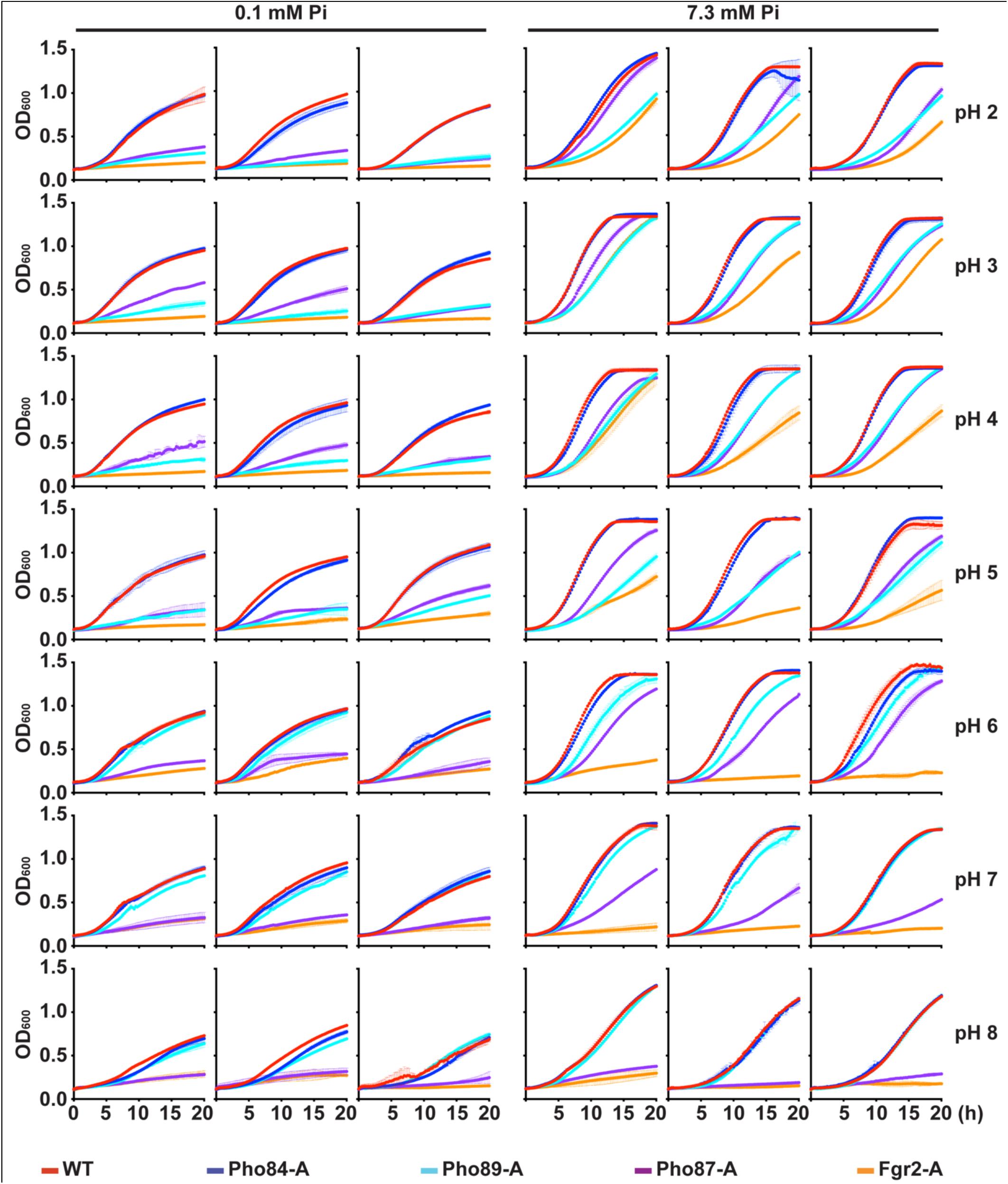
Individual growth curves summarized in Figure 4. Strains were grown as described in Fig. 4. Shown are 3 biological replicates performed on different days. Error bars SD of 2 or 3 technical replicates. Strains are WT (JKC915); Pho84-A: *pho87-/- pho89-/- fgr2-/- PHO84+/+* (JKC2788); Pho87-A: *pho84-/- pho89-/- fgr2-/- PHO87+/+* (JKC2777); Pho89-A: *pho84-/- pho87-/- fgr2-/- PHO89+/+* (JKC2783); Fgr2-A: *pho84-/- pho87-/- pho89-/- FGR2+/+* (JKC2758).

**S4 Fig.**
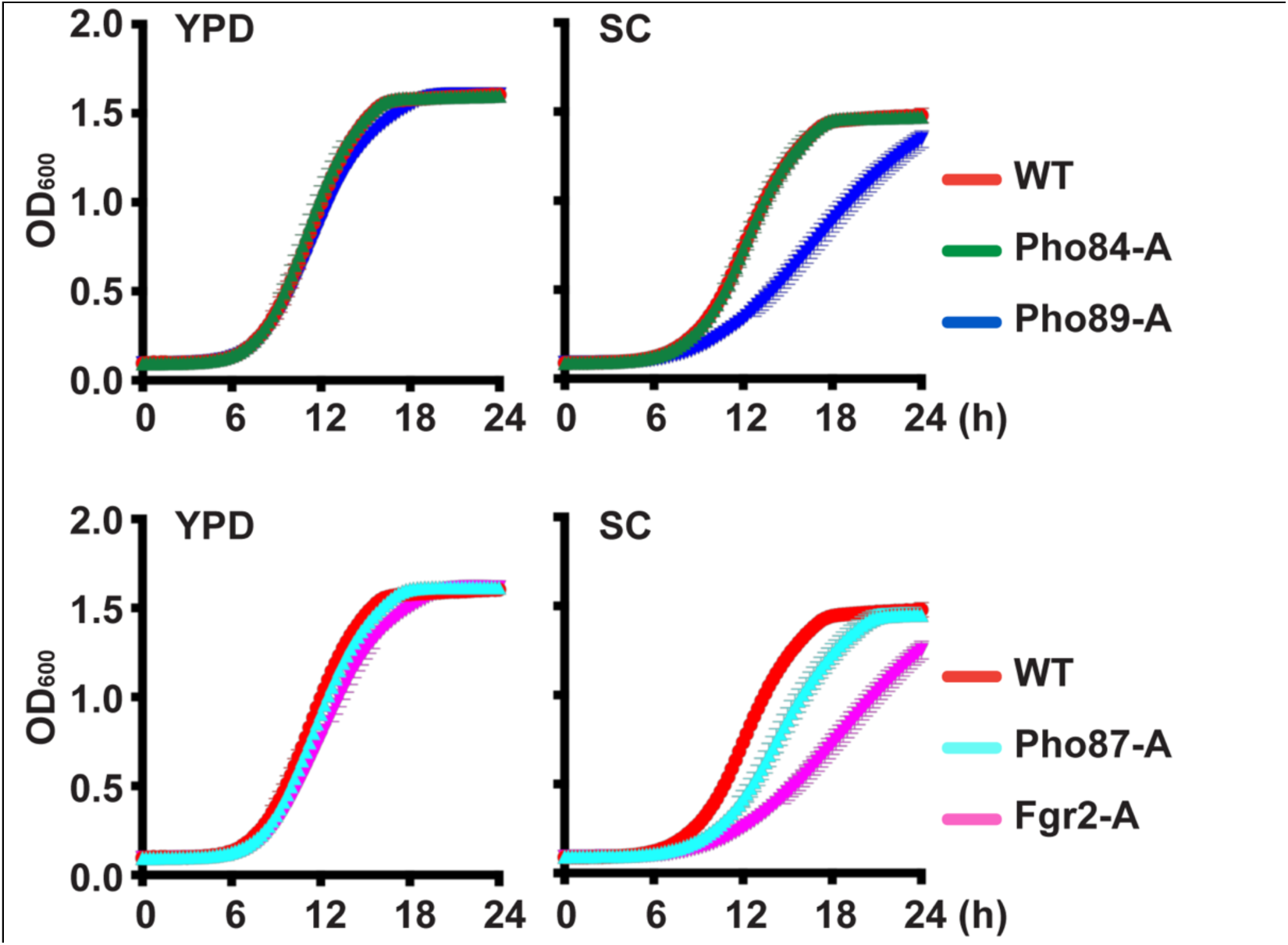
Pi transporter triple mutants had no growth defects in rich complex medium. Cells of indicated triple mutant genotypes were grown in YPD (left) and SC (right) and OD_600_ was monitored. Upper panels: strains expressing only one of 2 high-affinity transporters. Lower panels: Strains expressing only one of 2 low-affinity transporters. Pho84-A: *pho87-/- pho89-/- fgr2-/- PHO84+/+* (JKC2788); Pho89-A: *pho84-/- pho87-/- fgr2-/- PHO89+/+* (JKC2783); Pho87-A: *pho84-/- pho89-/- fgr2-/- PHO87+/+* (JKC2777); Fgr2-A: *pho84-/- pho87-/- pho89-/- FGR2+/+* (JKC2758). Representative of 3 biological replicates; error bars SD of 3 technical replicates.

**S5 Fig.**
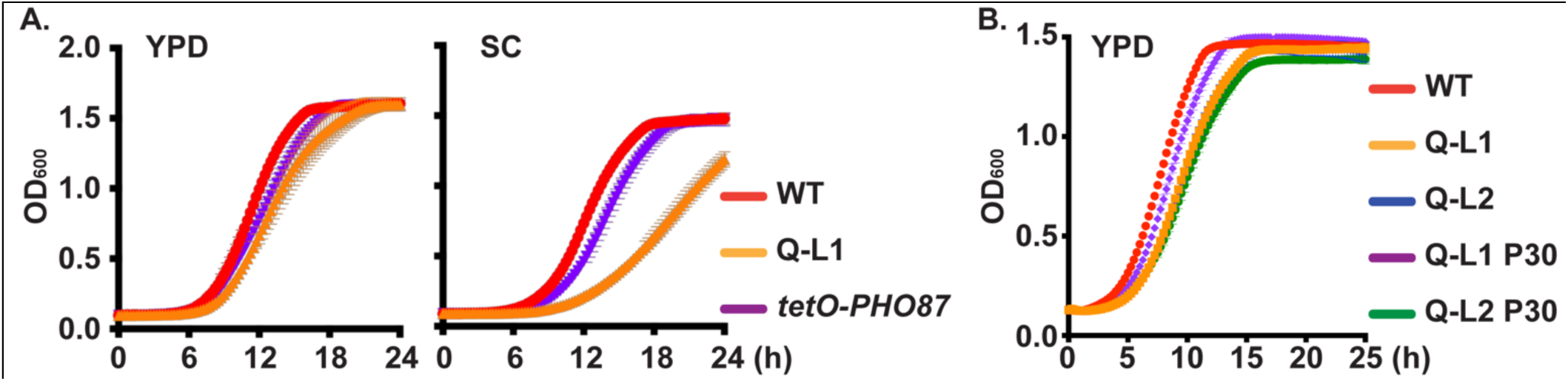
Cells whose single Pi transporter was expressed from *tetO*, as well as Q- cells and their Pi scarcity- evolved descendant populations had no substantial growth defects in rich complex medium. Strains were grown as in S4 Fig. **A.** Cells in which a single allele of one Pi transporter, *PHO87*, is expressed from repressible *tetO*, were grown in YPD and SC without doxycycline and compared with WT and Q- cells. WT (JKC915), Q- L1 (*pho84-/- pho89-/- pho87-/- fgr2-/-*, JKC2830), *tetO-PHO87* (*tetO-PHO87/pho87 pho84-/- pho89-/- fgr2-/- git2-4-/-*, JKC2969). **B.** Growth in YPD of WT (JKC915); Q- L1 (JKC2830) and Q- L2 (*pho84-/- pho89-/- pho87-/- fgr2-/-*, JKC2860). P30: population from the 30^th^ Pi scarcity passage.

