## Supplemental Tables 2-5 for "*Candida albicans’* inorganic phosphate transport and evolutionary adaptation to phosphate scarcity"

### S2 Table. Strains used in this study.

| *C. albicans* strain name | Parent | Genotype | Strain construction | Reference |
|---|---|---|---|---|
| SC5314 | | Wild type | | [1] |
| SN95 | | *arg4Δ/arg4Δ his1Δ/his1Δ IRO1/iro1Δ::λimm*$^{434}$ *URA3/ura3Δ::λimm*$^{434}$ | | [2] |
| JKC915 | SC5314 | *HIS1/his1::tetR-FRT* | | [3] |
| JKC1450 | JKC1423 | *pho84::HIS1/pho84::ARG4 his1/his1::tetR-FRT arg4/arg4 IRO1/iro1Δ::λimm*$^{434}$ *URA3/ura3Δ::λimm*$^{434}$ | | [4] |
| JKC2579 | JKC2566 | *pho84::HIS1/pho84::ARG4 pho89::uPAM*-FRT/pho89::uPAM-FRT-FLP-NAT1 his1/his1::tetR-FRT arg4/arg4 IRO1/iro1Δ::λimm*$^{434}$ *URA3/ura3Δ::λimm*$^{434}$ | JKC2566 transformed with KpnI/BsiWI digested pJK1481 to delete the 2$^{nd}$ allele of *PHO89*. | This work |
| JKC2592 | JKC2579 | *pho84::HIS1/pho84::ARG4 pho89::uPAM-FRT/pho89::uPAM-FRT his1/his1::tetR-FRT arg4/arg4 IRO1/iro1Δ::λimm*$^{434}$ *URA3/ura3Δ::λimm*$^{434}$ | JKC2579 *NAT1* flipped out | This work |
| JKC2536 | JKC915 | *PHO87/pho87::uPAM-FRT-FLP-NAT1 HIS1/his1::tetR-FRT* | JKC915 transformed with KpnI/BsiWI digested pJK1372 to delete the 1$^{st}$ allele of *PHO87*. | This work |
| JKC2728 | JKC2592 | *pho84::HIS1/pho84::ARG4 pho89::uPAM-FRT/pho89::uPAM-FRT FGR2/fgr2::uPAM-FRT-FLP-NAT1 his1/his1::tetR-FRT arg4/arg4 IRO1/iro1Δ::λimm*$^{434}$ *URA3/ura3Δ::λimm*$^{434}$ | JKC2592 transformed with KpnI/BsiWI digested pJK1485 to delete the 1$^{st}$ allele of *FGR2*. | This work |
| JKC2548 | JKC2536 | *PHO87/pho87::uPAM-FRT HIS1/his1::tetR-FRT* | JKC2536 *NAT1* flipped out | This work |
| JKC2737 | JKC2728 | *pho84::HIS1/pho84::ARG4 pho89::uPAM-FRT/pho89::uPAM-FRT FGR2/fgr2::uPAM-FRT his1/his1::tetR-FRT arg4/arg4 IRO1/iro1Δ::λimm*$^{434}$ *URA3/ura3Δ::λimm*$^{434}$ | JKC2728 *NAT1* flipped out | This work |
| JKC2573 | JKC2548 | *pho87::uPAM-FRT/pho87::uPAM-FRT-FLP-NAT1 HIS1/his1::tetR-FRT* | JKC2548 transformed with KpnI/BsiWI digested pJK1479 to delete the 2$^{nd}$ allele of *PHO87*. | This work |
| JKC2596 | JKC2554 | *pho84::HIS1/pho84::ARG4 pho87::uPAM-FRT/pho87::uPAM-FRT-FLP-NAT1 his1/his1::tetR-FRT arg4/arg4 IRO1/iro1Δ::λimm*$^{434}$ *URA3/ura3Δ::λimm*$^{434}$ | JKC2554 transformed with KpnI/BsiWI digested pJK1479 to delete the 2$^{nd}$ allele of *PHO87*. | This work |
| JKC2764 | JKC2737 | *pho84::HIS1/pho84::ARG4 pho89::uPAM-FRT/pho89::uPAM-FRT fgr2::uPAM-FRT/fgr2::uPAM-FRT-FLP-NAT1 his1/his1::tetR-FRT arg4/arg4 IRO1/iro1Δ::λimm*$^{434}$ *URA3/ura3Δ::λimm*$^{434}$ | JKC2737 transformed with KpnI/BsiWI digested pJK1488 to delete the 2$^{nd}$ allele of *FGR2*. | This work |
| JKC2581 | JKC2573 | *pho87::uPAM-FRT/pho87::uPAM-FRT HIS1/his1::tetR-FRT* | JKC2573 *NAT1* flipped out | This work |
| JKC2599 | JKC2596 | *pho84::HIS1/pho84::ARG4 pho87::uPAM-FRT/pho87::uPAM-FRT his1/his1::tetR-FRT arg4/arg4 IRO1/iro1Δ::λimm*$^{434}$ *URA3/ura3Δ::λimm*$^{434}$ | JKC2596 *NAT1* flipped out | This work |
| JKC2773 | JKC2764 | *pho84::HIS1/pho84::ARG4 pho89::uPAM-FRT/pho89::uPAM-FRT fgr2::uPAM-FRT/fgr2::uPAM-FRT his1/his1::tetR-FRT arg4/arg4 IRO1/iro1Δ::λimm*$^{434}$ *URA3/ura3Δ::λimm*$^{434}$ | JKC2764 *NAT1* flipped out | This work |
| JKC2638 | JKC2581 | *pho87::uPAM-FRT/pho87::uPAM-FRT PHO89/pho89::uPAM-FRT-FLP-NAT1 HIS1/his1::tetR-FRT* | JKC2581 transformed with KpnI/BsiWI digested pJK1384 to delete the 1$^{st}$ allele of *PHO89*. | This work |
| JKC2545 | JKC1450 | *pho84::HIS1/pho84::ARG4 PHO89/pho89::uPAM-FRT-FLP-NAT1 his1/his1::tetR-FRT arg4/arg4 IRO1/iro1Δ::λimm*$^{434}$ *URA3/ura3Δ::λimm*$^{434}$ | JKC1450 transformed with KpnI/BsiWI digested pJK1384 to delete the 1$^{st}$ allele of *PHO89*. | This work |
| JKC2539 | JKC1450 | *pho84::HIS1/pho84::ARG4 PHO87/pho87::uPAM-FRT-FLP-NAT1 his1/his1::tetR-FRT arg4/arg4 IRO1/iro1Δ::λimm*$^{434}$ *URA3/ura3Δ::λimm*$^{434}$ | JKC1450 transformed with KpnI/BsiWI digested pJK1372 to delete the 1$^{st}$ allele of *PHO87*. | This work |
| JKC2712 | JKC2599 | *pho84::HIS1/pho84::ARG4 pho87::uPAM-FRT/pho87::uPAM-FRT PHO89/pho89::uPAM-FRT-FLP-NAT1 his1/his1::tetR-FRT arg4/arg4 IRO1/iro1Δ::λimm*$^{434}$ | JKC2599 transformed with KpnI/BsiWI digested pJK1384 to delete the 1$^{st}$ allele of *PHO89*. | This work |

| | | | | |
|---|---|---|---|---|
| | | *URA3/ura3Δ::λimm*$^{434}$ | | |
| JKC2790 | JKC2773 | *pho84::HIS1/pho84::ARG4*<br>*pho89::uPAM-FRT/pho89::uPAM-FRT*<br>*fgr2::uPAM-FRT/fgr2::uPAM-FRT*<br>*PHO87/pho87::FRT-FLP-NAT1-tetO-PHO87*<br>*his1/his1::tetR-FRT arg4/arg4 IRO1/iro1Δ::λimm*$^{434}$<br>*URA3/ura3Δ::λimm*$^{434}$ | JKC2773 transformed with KpnI/NcoI digested pJK1375 to have one of the alleles of *PHO87* under the *tetO* control | This work |
| JKC2652 | JKC2638 | *pho87::uPAM-FRT/pho87::uPAM-FRT*<br>*PHO89/pho89::uPAM-FRT*<br>*HIS1/his1::tetR-FRT* | JKC2638 *NAT1* flipped out | This work |
| JKC2566 | JKC2545 | *pho84::HIS1/pho84::ARG4*<br>*PHO89/pho89::uPAM-FRT*<br>*his1/his1::tetR-FRT arg4/arg4 IRO1/iro1Δ::λimm*$^{434}$<br>*URA3/ura3Δ::λimm*$^{434}$ | JKC2545 *NAT1* flipped out | This work |
| JKC2554 | JKC2539 | *pho84::HIS1/pho84::ARG4*<br>*PHO87/pho87::uPAM-FRT*<br>*his1/his1::tetR-FRT arg4/arg4 IRO1/iro1Δ::λimm*$^{434}$<br>*URA3/ura3Δ::λimm*$^{434}$ | JKC2539 *NAT1* flipped out | This work |
| JKC2718 | JKC2712 | *pho84::HIS1/pho84::ARG4*<br>*pho87::uPAM-FRT/pho87::uPAM-FRT*<br>*PHO89/pho89::uPAM-FRT*<br>*his1/his1::tetR-FRT arg4/arg4 IRO1/iro1Δ::λimm*$^{434}$<br>*URA3/ura3Δ::λimm*$^{434}$ | JKC2712 *NAT1* flipped out | This work |
| JKC2793 | JKC2790 | *pho84::HIS1/pho84::ARG4*<br>*pho89::uPAM-FRT/pho89::uPAM-FRT*<br>*fgr2::uPAM-FRT/fgr2::uPAM-FRT*<br>*PHO87/pho87::FRT-tetO-PHO87*<br>*his1/his1::tetR-FRT arg4/arg4 IRO1/iro1Δ::λimm*$^{434}$<br>*URA3/ura3Δ::λimm*$^{434}$ | JKC2790 *NAT1* flipped out | This work |
| JKC2664 | JKC2652 | *pho87::uPAM-FRT/pho87::uPAM-FRT*<br>*pho89::uPAM-FRT/pho89::uPAM-FRT-FLP-NAT1*<br>*HIS1/his1::tetR-FRT* | JKC2652 transformed with KpnI/BsiWI digested pJK1481 to delete the 2$^{nd}$ allele of *PHO89*. | This work |
| JKC2755 | JKC2718 | *pho84::HIS1/pho84::ARG4*<br>*pho87::uPAM-FRT/pho87::uPAM-FRT*<br>*pho89::uPAM-FRT/pho89::uPAM-FRT-FLP-NAT1*<br>*his1/his1::tetR-FRT arg4/arg4 IRO1/iro1Δ::λimm*$^{434}$<br>*URA3/ura3Δ::λimm*$^{434}$ | JKC2718 transformed with KpnI/BsiWI digested pJK1481 to delete the 2$^{nd}$ allele of *PHO89*. | This work |
| JKC2799 | JKC2793 | *pho84::HIS1/pho84::ARG4*<br>*pho89::uPAM-FRT/pho89::uPAM-FRT*<br>*fgr2::uPAM-FRT/fgr2::uPAM-FRT*<br>*pho87::FRT-tetO-PHO87/pho87::uPAM-FRT-FLP-NAT1*<br>*his1/his1::tetR-FRT arg4/arg4 IRO1/iro1Δ::λimm*$^{434}$<br>*URA3/ura3Δ::λimm*$^{434}$ | JKC2793 transformed with KpnI/BsiWI digested pJK1372 to delete the wildtype allele of *PHO87*. | This work |
| JKC2800 | JKC2793 | Same as JKC2799, different isolate | Same as JKC2799, different isolate | This work |
| JKC2679 | JKC2664 | *pho87::uPAM-FRT/pho87::uPAM-FRT*<br>*pho89::uPAM-FRT/pho89::uPAM-FRT*<br>*HIS1/his1::tetR-FRT* | JKC2664 *NAT1* flipped out | This work |
| JKC2758 | JKC2755 | *pho84::HIS1/pho84::ARG4*<br>*pho87::uPAM-FRT/pho87::uPAM-FRT*<br>*pho89::uPAM-FRT/pho89::uPAM-FRT*<br>*his1/his1::tetR-FRT arg4/arg4 IRO1/iro1Δ::λimm*$^{434}$<br>*URA3/ura3Δ::λimm*$^{434}$ | JKC2755 *NAT1* flipped out | This work |
| JKC2804 | JKC2799 | *pho84::HIS1/pho84::ARG4*<br>*pho89::uPAM-FRT/pho89::uPAM-FRT*<br>*fgr2::uPAM-FRT/fgr2::uPAM-FRT*<br>*pho87::FRT-tetO-PHO87/pho87::uPAM-FRT*<br>*his1/his1::tetR-FRT arg4/arg4 IRO1/iro1Δ::λimm*$^{434}$<br>*URA3/ura3Δ::λimm*$^{434}$ | JKC2799 *NAT1* flipped out | This work |
| JKC2806 | JKC2800 | Same as JKC2804, flip-out from a different isolate | JKC2800 *NAT1* flipped out | This work |
| JKC2542 | JKC915 | *PHO89/pho89::uPAM-FRT-FLP-NAT1*<br>*HIS1/his1::tetR-FRT* | JKC915 transformed with KpnI/BsiWI digested pJK1384 to delete the 1$^{st}$ allele of *PHO89*. | This work |
| JKC2632 | JKC915 | *FGR2/fgr2::uPAM-FRT-FLP-NAT1*<br>*HIS1/his1::tetR-FRT* | JKC915 transformed with KpnI/BsiWI digested pJK1485 to delete the 1$^{st}$ allele of *FGR2*. | This work |
| JKC2734 | JKC2679 | *pho87::uPAM-FRT/pho87::uPAM-FRT*<br>*pho89::uPAM-FRT/pho89::uPAM-FRT*<br>*FGR2/fgr2::uPAM-FRT-FLP-NAT1*<br>*HIS1/his1::tetR-FRT* | JKC2679 transformed with KpnI/BsiWI digested pJK1485 to delete the 1$^{st}$ allele of *FGR2*. | This work |
| JKC2809 | JKC2758 | *pho84::HIS1/pho84::ARG4*<br>*pho87::uPAM-FRT/pho87::uPAM-FRT*<br>*pho89::uPAM-FRT/pho89::uPAM-FRT*<br>*FGR2/fgr2::uPAM-FRT-FLP-NAT1* | JKC2758 transformed with KpnI/BsiWI digested pJK1485 to delete the 1$^{st}$ allele of *FGR2*. | This work |

| | | | | |
|---|---|---|---|---|
| | | his1/his1::tetR-FRT arg4/arg4 IRO1/iro1Δ::λimm$^{434}$<br>URA3/ura3Δ::λimm$^{434}$ | | |
| JKC2915 | JKC2804 | pho84::HIS1/pho84::ARG4<br>pho89::uPAM-FRT/pho89::uPAM-FRT<br>fgr2::uPAM-FRT/fgr2::uPAM-FRT<br>pho87::FRT-tetO-PHO87/pho87::uPAM-FRT<br>GIT2-4/git2-4::uPAM-FRT-FLP-NAT1<br>his1/his1::tetR-FRT arg4/arg4 IRO1/iro1Δ::λimm$^{434}$<br>URA3/ura3Δ::λimm$^{434}$ | JKC2804 transformed with KpnI/BsiWI digested pJK1543 to delete the 1st allele of GIT2-4. | This work |
| JKC2917 | JKC2806 | Same as JKC2915, derived from JKC2806. | JKC2806 transformed with KpnI/BsiWI digested pJK1543 to delete the 1st allele of GIT2-4. | This work |
| JKC2560 | JKC2542 | PHO89/pho89::uPAM-FRT<br>HIS1/his1::tetR-FRT | JKC2542 NAT1 flipped out | This work |
| JKC2641 | JKC2632 | FGR2/fgr2::uPAM-FRT<br>HIS1/his1::tetR-FRT | JKC2632 NAT1 flipped out | This work |
| JKC2749 | JKC2734 | pho87::uPAM-FRT/pho87::uPAM-FRT<br>pho89::uPAM-FRT/pho89::uPAM-FRT<br>FGR2/fgr2::uPAM-FRT<br>HIS1/his1::tetR-FRT | JKC2734 NAT1 flipped out | This work |
| JKC2812 | JKC2809 | pho84::HIS1/pho84::ARG4<br>pho87::uPAM-FRT/pho87::uPAM-FRT<br>pho89::uPAM-FRT/pho89::uPAM-FRT<br>FGR2/fgr2::uPAM-FRT<br>his1/his1::tetR-FRT arg4/arg4 IRO1/iro1Δ::λimm$^{434}$<br>URA3/ura3Δ::λimm$^{434}$ | JKC2809 NAT1 flipped out | This work |
| JKC2826 | JKC2812 | pho84::HIS1/pho84::ARG4<br>pho87::uPAM-FRT/pho87::uPAM-FRT<br>pho89::uPAM-FRT/pho89::uPAM-FRT<br>fgr2::uPAM-FRT/fgr2::uPAM-FRT-FLP-NAT1<br>his1/his1::tetR-FRT arg4/arg4 IRO1/iro1Δ::λimm$^{434}$<br>URA3/ura3Δ::λimm$^{434}$ | JKC2812 transformed with KpnI/BsiWI digested pJK1488 to delete the 2nd allele of FGR2. | This work |
| JKC2926 | JKC2915 | pho84::HIS1/pho84::ARG4<br>pho89::uPAM-FRT/pho89::uPAM-FRT<br>fgr2::uPAM-FRT/fgr2::uPAM-FRT<br>pho87::FRT-tetO-PHO87/pho87::uPAM-FRT<br>GIT2-4/git2-4::uPAM-FRT<br>his1/his1::tetR-FRT arg4/arg4 IRO1/iro1Δ::λimm$^{434}$<br>URA3/ura3Δ::λimm$^{434}$ | JKC2915 NAT1 flipped out | This work |
| JKC2930 | JKC2917 | Same as JKC2926, derived from JKC2806. | JKC2917 NAT1 flipped out | This work |
| JKC2575 | JKC2560 | pho89::uPAM-FRT/pho89::uPAM-FRT-FLP-NAT1<br>HIS1/his1::tetR-FRT | JKC2560 transformed with KpnI/BsiWI digested pJK1481 to delete the 2nd allele of PHO89. | This work |
| JKC2658 | JKC2641 | fgr2::uPAM-FRT/fgr2::uPAM-FRT-FLP-NAT1<br>HIS1/his1::tetR-FRT | JKC2641 transformed with KpnI/BsiWI digested pJK1488 to delete the 2nd allele of FGR2. | This work |
| JKC2772 | JKC2749 | pho87::uPAM-FRT/pho87::uPAM-FRT<br>pho89::uPAM-FRT/pho89::uPAM-FRT<br>fgr2::uPAM-FRT/fgr2::uPAM-FRT-FLP-NAT1<br>HIS1/his1::tetR-FRT | JKC2749 transformed with KpnI/BsiWI digested pJK1488 to delete the 2nd allele of FGR2. | This work |
| JKC2766 | JKC2737 | pho84::HIS1/pho84::ARG4<br>pho89::uPAM-FRT/pho89::uPAM-FRT<br>fgr2::uPAM-FRT/fgr2::uPAM-FRT-FLP-NAT1<br>his1/his1::tetR-FRT arg4/arg4 IRO1/iro1Δ::λimm$^{434}$<br>URA3/ura3Δ::λimm$^{434}$ | JKC2737 transformed with KpnI/BsiWI digested pJK1488 to delete the 2nd allele of FGR2. | This work |
| JKC2769 | JKC2743 | pho84::HIS1/pho84::ARG4<br>pho87::uPAM-FRT/pho87::uPAM-FRT<br>fgr2::uPAM-FRT/fgr2::uPAM-FRT-FLP-NAT1<br>his1/his1::tetR-FRT arg4/arg4 IRO1/iro1Δ::λimm$^{434}$<br>URA3/ura3Δ::λimm$^{434}$ | JKC2743 transformed with KpnI/BsiWI digested pJK1488 to delete the 2nd allele of FGR2. | This work |
| JKC2743 | JKC2731 | pho84::HIS1/pho84::ARG4<br>pho87::uPAM-FRT/pho87::uPAM-FRT<br>FGR2/fgr2::uPAM-FRT<br>his1/his1::tetR-FRT arg4/arg4 IRO1/iro1Δ::λimm$^{434}$<br>URA3/ura3Δ::λimm$^{434}$ | JKC2731 NAT1 flipped out | This work |
| JKC2731 | JKC2599 | pho84::HIS1/pho84::ARG4<br>pho87::uPAM-FRT/pho87::uPAM-FRT<br>FGR2/fgr2::uPAM-FRT-FLP-NAT1<br>his1/his1::tetR-FRT arg4/arg4 IRO1/iro1Δ::λimm$^{434}$<br>URA3/ura3Δ::λimm$^{434}$ | JKC2599 transformed with KpnI/BsiWI digested pJK1485 to delete the 1st allele of FGR2. | This work |
| JKC2844 | JKC2812 | Same as JKC2826. | Same as JKC2826, different isolate. | This work |
| JKC2957 | JKC2926 | pho84::HIS1/pho84::ARG4<br>pho89::uPAM-FRT/pho89::uPAM-FRT | JKC2926 transformed with KpnI/BsiWI digested pJK1545 to | This work |

| | | | | |
|---|---|---|---|---|
| | | *fgr2::uPAM-FRT/fgr2::uPAM-FRT*<br>*pho87::FRT-tetO-PHO87/pho87::uPAM-FRT*<br>*git2-4::uPAM-FRT/git2-4::uPAM-FRT-FLP-NAT1*<br>*his1/his1::tetR-FRT arg4/arg4 IRO1/iro1Δ::λimm*$^{434}$<br>*URA3/ura3Δ::λimm*$^{434}$ | delete the 2$^{nd}$ allele of *GIT2-4*. | |
| JKC2958 | JKC2926 | Same as JKC2957. | Same as JKC2957, different isolate. | This work |
| JKC2961 | JKC2930 | Same as JKC2957, derived from JKC2930 | Same as JKC2957, derived from JKC2930 | This work |
| JKC2585<br>"*pho89-/-*" | JKC2575 | *pho89::uPAM-FRT/pho89::uPAM-FRT*<br>*HIS1/his1::tetR-FRT* | JKC2575 *NAT1* flipped out | This work |
| JKC2667<br>"*fgr2-/-*" | JKC2658 | *fgr2::uPAM-FRT/fgr2::uPAM-FRT*<br>*HIS1/his1::tetR-FRT* | JKC2658 *NAT1* flipped out | This work |
| JKC2788<br>"Pho84-A" | JKC2772 | *pho87::uPAM-FRT/pho87::uPAM-FRT*<br>*pho89::uPAM-FRT/pho89::uPAM-FRT*<br>*fgr2::uPAM-FRT/fgr2::uPAM-FRT*<br>*HIS1/his1::tetR-FRT* | JKC2772 *NAT1* flipped out | This work |
| JKC2777<br>"Pho87-A" | JKC2766 | *pho84::HIS1/pho84::ARG4*<br>*pho89::uPAM-FRT/pho89::uPAM-FRT*<br>*fgr2::uPAM-FRT/fgr2::uPAM-FRT*<br>*his1/his1::tetR-FRT arg4/arg4 IRO1/iro1Δ::λimm*$^{434}$<br>*URA3/ura3Δ::λimm*$^{434}$ | JKC2766 *NAT1* flipped out | This work |
| JKC2783<br>"Pho89-A" | JKC2769 | *pho84::HIS1/pho84::ARG4*<br>*pho87::uPAM-FRT/pho87::uPAM-FRT*<br>*fgr2::uPAM-FRT/fgr2::uPAM-FRT*<br>*his1/his1::tetR-FRT arg4/arg4 IRO1/iro1Δ::λimm*$^{434}$<br>*URA3/ura3Δ::λimm*$^{434}$ | JKC2769 *NAT1* flipped out | This work |
| JKC2830<br>"Q-L1" | JKC2826 | *pho84::HIS1/pho84::ARG4*<br>*pho87::uPAM-FRT/pho87::uPAM-FRT*<br>*pho89::uPAM-FRT/pho89::uPAM-FRT*<br>*fgr2::uPAM-FRT/fgr2::uPAM-FRT*<br>*his1/his1::tetR-FRT arg4/arg4 IRO1/iro1Δ::λimm*$^{434}$<br>*URA3/ura3Δ::λimm*$^{434}$ | JKC2826 *NAT1* flipped out | This work |
| JKC2831<br>"Q-" | JKC2826 | Same as JKC2830, different flip-out isolate | JKC2826 *NAT1* flipped out | This work |
| JKC2858<br>"Q-" | JKC2844 | Same as JKC2830, flip-out from JKC2844 | JKC2844 *NAT1* flipped out | This work |
| JKC2859<br>"Q-" | JKC2844 | Same as JKC2830, flip-out from JKC2844<br>Different isolate from JKC2858 | JKC2844 *NAT1* flipped out | This work |
| JKC2845 | JKC2812 | Same as JKC2826. | Same as JKC2826, different isolate. | This work |
| JKC2860<br>"Q-L2" | JKC2845 | Same as JKC2830, flip-out from JKC2845 | JKC2845 *NAT1* flipped out | This work |
| JKC2861<br>"Q-" | JKC2845 | Same as JKC2830, flip-out from JKC2845 | JKC2845 *NAT1* flipped out | This work |
| JKC2967<br>"Septuple mutant *tetO-PHO87*" | JKC2957 | *pho84::HIS1/pho84::ARG4*<br>*pho89::uPAM-FRT/pho89::uPAM-FRT*<br>*fgr2::uPAM-FRT/fgr2::uPAM-FRT*<br>*pho87::FRT-tetO-PHO87/pho87::uPAM-FRT*<br>*git2-4::uPAM-FRT/git2-4::uPAM-FRT*<br>*his1/his1::tetR-FRT arg4/arg4 IRO1/iro1Δ::λimm*$^{434}$<br>*URA3/ura3Δ::λimm*$^{434}$ | JKC2957 *NAT1* flipped out | This work |
| JKC2969<br>"Septuple mutant *tetO-PHO87*" | JKC2958 | Same as JKC2967, flip-out from JKC2958. | JKC2958 *NAT1* flipped out | This work |
| JKC2973<br>"Septuple mutant *tetO-PHO87*" | JKC2961 | Same as JKC2967, derived from JKC2930 | JKC2961 *NAT1* flipped out | This work |

**uPAM* is an artificially designed sequence (universal-PAM sequence) in our *FLP-NAT1* cassette that we use in other work for CRISPR guide RNA recognition; not used in this work.

#### Construction of strains with multiple mutations

Construction of the Pho84-A strain (*pho87-/- pho89-/- fgr2-/-* triple mutant):

In brief, Pho84-Alone (Pho84-A) strain JKC2788 was constructed as follows: wild type strain JKC915 [3] was transformed with pJK1372 and pJK1479 to sequentially delete two alleles of *PHO87*, resulting in JKC2581 (*pho87-/-*). JKC2581 (*pho87-/-*) was then transformed with pJK1384 and pJK1481 to sequentially delete two alleles of *PHO89*, resulting in JKC2679 (*pho87-*

*/- pho89-/-*). JKC2679 (*pho87-/- pho89-/-*) was eventually transformed with pJK1485 and pJK1488 to sequentially delete two alleles of *FGR2*, resulting in JKC2788 (*pho87-/- pho89-/- fgr2-/-*).

<u>Construction of the Pho87-A strain (*pho84-/- pho89-/- fgr2-/-* triple mutant):</u>

In brief, Pho87-Alone (Pho87-A) strain JKC2777 was constructed as follows: JKC1450 (*pho84-/-*) [4] was transformed with pJK1384 and pJK1481 to sequentially delete two alleles of *PHO89*, resulting in JKC2592 (*pho84-/- pho89-/-*). JKC2592 (*pho84-/- pho89-/-*) was eventually transformed with pJK1485 and pJK1488 to sequentially delete two alleles of *FGR2*, resulting in JKC2777 (*pho84-/- pho89-/- fgr2-/-*).

<u>Construction of the Pho89-A strain (*pho84-/- pho87-/- fgr2-/-* triple mutant):</u>

In brief, Pho89-Alone (Pho89-A) strain JKC2783 was constructed as follows: JKC1450 (*pho84-/-*) [4] was transformed with pJK1372 and pJK1479 to sequentially delete two alleles of *PHO87*, resulting in JKC2599 (*pho84-/- pho87-/-*). JKC2599 (*pho84-/- pho87-/-*) was eventually transformed with pJK1485 and pJK1488 to sequentially delete two alleles of *FGR2*, resulting in JKC2783 (*pho84-/- pho87-/- fgr2-/-*).

<u>Construction of the Fgr2-A strain (*pho84-/- pho87-/- pho89-/-* triple mutant):</u>

In brief, Fgr2-Alone (Fgr2-A) strain JKC2758 was constructed as follows: JKC1450 (*pho84-/-*) [4] was transformed with pJK1372 and pJK1479 to sequentially delete two alleles of *PHO87*, resulting in JKC2599 (*pho84-/- pho87-/-*). JKC2599 (*pho84-/- pho87-/-*) was eventually transformed with pJK1384 and pJK1481 to sequentially delete two alleles of *PHO89*, resulting in JKC2758 (*pho84-/- pho87-/- pho89-/-*).

<u>Construction of quadruple mutants:</u>

In brief, using quadruple mutant strain JKC2830 as an example, JKC1450 (*pho84-/-*) [4] was transformed with pJK1372 and pJK1479 to sequentially delete two alleles of *PHO87*, resulting in JKC2599 (*pho84-/- pho87-/-*). JKC2599 (*pho84-/- pho87-/-*) was then transformed with pJK1384 and pJK1481 to sequentially delete two alleles of *PHO89*, resulting in JKC2758 (*pho84-/- pho87-/- pho89-/-*). JKC2758 (*pho84-/- pho87-/- pho89-/-*) was eventually transformed with pJK1485 and pJK1488 to sequentially delete two alleles of *FGR2*, resulting in JKC2830 (*pho84-/- pho87-/- pho89-/- fgr2-/-*).

<u>Construction of septuple mutants:</u>

In brief, using septuple mutant strain JKC2969 as an example, JKC1450 (*pho84-/-*) [4] was transformed with pJK1384 and pJK1481 to sequentially delete two alleles of *PHO89*, resulting in JKC2592 (*pho84-/- pho89-/-*). JKC2592 (*pho84-/- pho89-/-*) was then transformed with pJK1485 and pJK1488 to sequentially delete two alleles of *FGR2*, resulting in JKC2773 (*pho84-/- pho89-/- fgr2-/-*). JKC2773 (*pho84-/- pho89-/- fgr2-/-*) was first transformed with pJK1375 to have one allele of *PHO87* under *tetO* control, then transformed with pJK1372 to delete the WT allele of *PHO87*, resulting in JKC2804 (*pho84-/- pho89-/- fgr2-/- pho87/tetO-PHO87*). JKC2804 (*pho84-/- pho89-/- fgr2-/- pho87/tetO-PHO87*) was eventually transformed with pJK1543 and pJK1545 to sequentially delete two alleles of *GIT2-4*, resulting in JKC2969 (*pho84-/- pho89-/- fgr2-/- PHO87-/tetO git2-4-/-*).

### S3 Table. Plasmids used in this study.

| Plasmid | Description | Source (Reference) |
|---|---|---|
| pJK1000 | *FLP-NAT1 tetO-PES1* construct, vector backbone is pLitmus28 (New England Biolabs) | [3] |
| pJK1372 | *FLP-NAT1 pho87* deletion construct, derived from pJK1364. Product of fjk1846 and r1862 using SC5314 genomic DNA as template was ligated into pJK1364 using KpnI/ApaI sites. | [5] |
| pJK1375 | *FLP-NAT1 tetO-PHO87* construct, derived from pJK1000. Product of fjk1854 and rjk1855 using SC5314 genomic DNA as template and product of fjk1856 and rjk1857 using SC5314 genomic DNA as template were ligated into pJK1000 using KpnI/ApaI and SacII/NcoI sites, respectively. | This work |
| pJK1384 | *FLP-NAT1 pho89* deletion construct, derived from pJK1372. Product of fjk1869 and rjk1870 using SC5314 genomic DNA as template and product of fjk1871 and rjk1872 using SC5314 genomic DNA as template were ligated into pJK1372 using KpnI/AscI and NotI/BsiWI sites, respectively. | This work |
| pJK1479 | *FLP-NAT1 pho87* 2nd allele deletion construct, derived from pJK1372. Product of fjk2032 and rjk2033 using SC5314 genomic DNA as template was ligated into pJK1372 using KpnI/AscI sites. | This work |
| pJK1481 | *FLP-NAT1 pho89* 2nd allele deletion construct, derived from pJK1372. Product of fjk2034 and rjk2035 using SC5314 genomic DNA as template and product of fjk1871 and rjk1872 using SC5314 genomic DNA as template were ligated into pJK1372 using KpnI/AscI and NotI/BsiWI sites, respectively. | This work |
| pJK1485 | *FLP-NAT1 fgr2* deletion construct, derived from pJK1372. Product of fjk2037 and rjk2038 using SC5314 genomic DNA as template and product of fjk2041 and rjk2042 using SC5314 genomic DNA as template were ligated into pJK1372 using KpnI/AscI and NotI/BsiWI sites, respectively. | This work |
| pJK1488 | *FLP-NAT1 fgr2* 2nd allele deletion construct, derived from pJK1372. Product of fjk2039 and rjk2040 using SC5314 genomic DNA as template and product of fjk2041 and rjk2042 using SC5314 genomic DNA as template were ligated into pJK1372 using KpnI/AscI and NotI/BsiWI sites, respectively. | This work |
| pJK1543 | *FLP-NAT1 git2-4* deletion construct, derived from pJK1372. Product of fjk2197 and rjk2198 using SC5314 genomic DNA as template and product of fjk2201 and rjk2202 using SC5314 genomic DNA as template were ligated into pJK1372 using KpnI/AscI and NotI/BsiWI sites, respectively. | This work |
| pJK1545 | *FLP-NAT1 git2-4* 2nd allele deletion construct, derived from pJK1372. Product of fjk2199 and rjk2200 using SC5314 genomic DNA as template and product of fjk2201 and rjk2202 using SC5314 genomic DNA as template were ligated into pJK1372 using KpnI/AscI and NotI/BsiWI sites, respectively. | This work |

### S4 Table. Oligonucleotides used in this study.

| Primer name | Purpose | Sequence 5' to 3'<br>(lower cases - restriction enzyme recognition sites) |
|---|---|---|
| fjk1854 | Forward oligo to amplify the *tetO-PHO87* construct upstream homologous sequence<br>Also used to verify the 5'end of *pho87* deletion mutant | CATCCGggtaccCAATAGAGCGGGAATGGAAA |
| rjk1855 | Reverse oligo to amplify the *tetO-PHO87* construct upstream homologous sequence | GATCgggcccTCAATTTCACCCATCAAAAACA |
| fjk1856 | Forward oligo to amplify the *tetO-PHO87* construct downstream homologous sequence | GGATCCccgcggATGAAGTTTTCTCATTCATTG |
| rjk1857 | Reverse oligo to amplify the *tetO-PHO87* construct downstream homologous sequence | CTCATGccatggGATTTCTAAATCACTTTCACTACCATAA |
| fjk1877 | Forward oligo to verify the 5'end of *tetO-PHO87* integration | GCTTGCTGGTTTAACCTGAG |
| rjk1878 | Reverse oligo to verify the 3'end of *tetO-PHO87* integration | GTGTCATGTTCACCATTGGA |
| fjk1869 | Forward oligo to amplify the *pho89* deletion construct upstream homologous sequence | CATCCGggtaccTTGCAATTATTTTTCTTGTCCAAA |
| rjk1870 | Reverse oligo to amplify the *pho89* deletion construct upstream homologous sequence | GATggcgcgccTGTATATATTTGAATTTATTTGTTGTTG |
| fjk1871 | Forward oligo to amplify the *pho89* deletion construct downstream homologous sequence | AAGGTAAGCAgcggccgcACGTTGTTTGTGGTTTCAATTT<br>AG |
| rjk1872 | Reverse oligo to amplify the *pho89* deletion construct downstream homologous sequence | CTCATGcgtacgCCTTGGCATAGCATTAGTAATCA |
| fjk1873 | Forward oligo to verify the 5'end of *pho89* deletion mutant | CATCCGggtaccCGACAAACATGCTTCCTTGA |
| rjk1885 | Reverse oligo to verify the 3'end of *pho89* deletion mutant | CGACACTTCTTGGATTTGCT |

| | | |
|---|---|---|
| fjk2032 | Forward oligo to amplify the *pho87* deletion-2nd allele construct upstream homologous sequence | CATCCGggtaccATGAAGTTTTCTCATTCATTGAAATTTAATGC |
| rjk2033 | Reverse oligo to amplify the *pho87* deletion-2nd allele construct upstream homologous sequence | GATggcgcgccCATTGGTGTTGTTATTTCTTTTGAA |
| rjk1339 | Reverse oligo to verify the 5'end of *integration of 'FLP-NAT1'* cassette containing constructs | TGGTGTGTTGTTGACAGGCAAC |
| fjk490 | Forward oligo to verify the 3'end integration of '*FLP-NAT1*' cassette containing constructs | TCAAGGAGGGTATTCTGGGC |
| fjk1835 | Forward oligo to verify the 3'end integration of '*FLP-NAT1-tetO'* constructs | TGTCGTTTCTGATGGGCTTT |
| rjk1879 | Reverse oligo to verify the 3'end of *pho87* deletion mutant | AACAACAACCACAACCACAA |
| fjk2034 | Forward oligo to amplify the *pho89* deletion-2nd allele construct upstream homologous sequence | CATCCGggtaccATGGCTTTACATCAATTTGATTATTTGTTTG |
| rjk2035 | Reverse oligo to amplify the *pho89* deletion-2nd allele construct upstream homologous sequence | GATggcgcgccGTCATAGTCAACATCAACACAGCAG |
| fjk2037 | Forward oligo to amplify the *fgr2* deletion construct upstream homologous sequence | CATCAAggtaccAACTCTATTTCTCGAAGCTGTCAAA |
| rjk2038 | Reverse oligo to amplify the *fgr2* deletion construct upstream homologous sequence | GATggcgcgccAGACTCAATTACGAAACACAAGACC |
| fjk2039 | Forward oligo to amplify the *fgr2* deletion-2nd allele construct upstream homologous sequence | CATCAAggtaccTCTCACACGTTCCAAATAAGAAAAC |
| rjk2040 | Reverse oligo to amplify the *fgr2* deletion-2nd allele construct upstream homologous sequence | GATggcgcgccTTTCAGTTACAGACGGAATGAATAA |
| fjk2041 | Forward oligo to amplify the *fgr2* deletion construct downstream homologous sequence | TTGGTAAGCAgcggccgcATTTAATATCCAACTTAGCTCAAATAA |
| rjk2042 | Reverse oligo to amplify the *fgr2* deletion construct downstream homologous sequence | CTCATGcgtacgCCCTTTGAAGATATATTTGATGAAACC |
| fjk2043 | Forward oligo to verify the 5'end of *fgr2* deletion mutant | TATAACCTAGCAGAATAGCCGATTG |
| rjk2044 | Reverse oligo to verify the 3'end of *fgr2* deletion mutant | ACGTATTGGTTGAATTTTGGAGTAG |
| fjk2197 | Forward oligo to amplify the *git2-4* deletion construct upstream homologous sequence | CATCAAggtaccATTTAGGCTGCAAAAAGAGAAAAAT |
| rjk2198 | Reverse oligo to amplify the *git2-4* deletion construct upstream homologous sequence | GATggcgcgccTGCTCTGATTAATCTTCGACCTAGT |
| fjk2199 | Forward oligo to amplify the *git2-4* deletion-2nd allele construct upstream homologous sequence | CATCAAggtaccTCAATTAATAGTCTTTGCCATAAACA |
| rjk2200 | Reverse oligo to amplify the *git2-4* deletion-2nd allele construct upstream homologous sequence | GATggcgcgccCCATGTTATTATTTGTTTGACTTGTTAGG |
| fjk2201 | Forward oligo to amplify the *git2-4* deletion construct downstream homologous sequence | TTGGTAAGCAgcggccgcGGAGGTTCAACTTTGCAGGT |
| rjk2202 | Reverse oligo to amplify the *git2-4* deletion construct downstream homologous sequence | CTCATGcgtacgTTGCCGAAGTGGGTTTGTAT |
| fjk2203 | Forward oligo to verify the 5'end of *git2-4* deletion mutant | ATACACACACTCCCAAAAACTCATT |
| rjk2204 | Reverse oligo to verify the 3'end of *git2-4* deletion mutant | GCCACCAAGTAGGTTTGGAA |

### S5 Table. Antibodies used in this study.

| Purpose | Antigen recognized | Species | Source or Reference |
|---|---|---|---|
| loading control | tubulin | rat | Abcam, cat. # ab6161 |
| P-S6 | phospho (S/T)-Akt substrate | rabbit | Cell Signaling Technology, cat. # 9611 |
| secondary | rat IgG | goat | Santa Cruz Biotechnology, cat. #97057 |
| secondary | rabbit IgG | goat | Cell Signaling Technology, cat. #7074S |
